# A label-free method to track individuals and lineages of budding cells

**DOI:** 10.1101/2022.05.11.491488

**Authors:** Julian M. J. Pietsch, Alán F. Muñoz, Diane-Yayra A. Adjavon, Iseabail Farquhar, Ivan B. N. Clark, Peter S. Swain

## Abstract

Much of biochemical regulation ultimately controls growth rate, particularly in microbes. Although time-lapse microscopy visualises cells, determining their growth rates is challenging because cells often overlap in images, particularly for those that divide asymmetrically, like *Saccharomyces cerevisiae*. Here we present the Birth Annotator for Budding Yeast (BABY), an algorithm to determine single-cell growth rates from label-free images. Using a convolutional neural network, BABY resolves overlaps through separating cells by size and assigns buds to mothers by identifying bud necks. BABY uses machine learning to track cells and determine lineages, estimates growth rates as the rate of change of volumes, and identifies cytokinesis by how growth varies. Using BABY and a microfluidic device, we show that bud growth is first sizer- then timer-controlled, that the nuclear concentration of Sfp1, a regulator of ribosome biogenesis, varies before the growth rate does, and that growth rate can be used for real-time control. Growth rate and fitness are strongly correlated, and BABY should therefore generate much biological insight.

## Introduction

For microbes, growth rate correlates strongly with fitness [1]. Cells increase growth rates through balancing their synthesis of ribosomes with their intake of nutrients [2–4] and target a particular size through coordinating growth with division [5–8]. Multicellular organisms, too, not only coordinate growth over time but also in space to both size and position cells correctly [9].

To understand how growth rate is regulated, studying single cells is often most informative [10]. Time-lapse microscopy, particularly with microfluidic technology to control the extracellular environment [11,12], has been pivotal, allowing, for example, studies of the cell-cycle machinery [7], of the control of cell size [13–15], of antibiotic effects [16, 17], of the response to stress [18–20], of feedback between growth and metabolism [21], and of ageing [22].

For cells that bud, like *Saccharomyces cerevisiae*, estimating an instantaneous growth rate for individual cells is challenging. *S. cerevisiae* grows by forming a bud that increases in size while the volume of the rest of the cell remains relatively unchanged. Although growth rate is typically reported as the rate of change of volume [13, 15, 23–26], which approximates a cell’s increase in mass, these estimates rely on solving multiple computational challenges: accurately determining the outlines of cells – particularly buds – in an image, extrapolating these outlines to volumes, tracking cells over time, assigning buds to the appropriate mother cells, and identifying births. Growth rates for budding yeast are therefore often only reported for isolated cells using low-throughput and semi-automated methods [13, 26]. In contrast, for rod-shaped cells that divide symmetrically, like *Escherichia coli*, the growth rate can be found more simply, as the rate of change of a cell’s length [21].

A particular difficulty is identifying cell boundaries because neighbouring cells in images often overlap – yeast, like other microbes, grows in colonies. Although software that automatically identifies and tracks cells in bright-field and phase-contrast images is well established [27–31] – with deep learning now improving both accuracy and speed [32–34], only a few algorithms allow for overlaps [30,33]. For example, the convolutional neural network U-net [35], a workhorse in biomedical image processing, performs semantic segmentation, lumping adjacent objects in the same category under one label, and instances of individual cells must be found using post-processing. Even then different instances typically cannot overlap [32, 34]. Other deep-learning approaches, like Mask-RCNN [36] and extended U-nets like StarDist [37], can identify overlapping instances in principle, but typically do not, by either implementation [37] or the labelling of the training data [33]. Furthermore, assigning lineages and births is often performed manually [13, 23] or through fluorescent markers [15, 25], but such markers use an imaging channel.

Here we describe the Birth Annotator for Budding Yeast (BABY), a complete pipeline to robustly and accurately determine single-cell growth rates from label-free images of budding yeast. In developing BABY, we solved multiple image-processing challenges generated by cells dividing asymmetrically. BABY resolves overlapping instances – buds, particularly small ones, usually overlap with their mothers and neighbouring cells – by extending the U-net architecture with custom training targets and post processing. It tracks cells between time points with a machine-learning algorithm, which is able to resolve any large movements of cells from one image to the next, and assigns lineages with another informed by the U-net. These innovations substantially improve performance. BABY produces high-fidelity time series of the volumes of both mother cells and buds and so the instantaneous growth rates of single cells at fine temporal resolution.

Using BABY, we find that concurrent behaviour in the growth rates of the mother cell and its bud enables us to identify cytokinesis from bright-field images and so accurately estimate birth times. We see a peak in growth rate during the S/G2/M phase of the cell cycle and show that this peak marks a transition in the bud’s growth from being sizer- to timer-controlled. Studying Sfp1, a regulator of ribosome synthesis, we observe correlated fluctuations between this regulator’s nuclear concentration and growth rate. Finally, we demonstrate that BABY is sufficiently fast to enable real-time control, running an experiment where changes in the extracellular medium are triggered when the growth of the cells being imaged crosses a pre-determined threshold.

## Results

### Segmenting overlapping cells using a multi-target convolutional neural network

To estimate single-cell growth rates from time-lapse microscopy images, accurately identifying and segmenting cells is essential. Poorly defined outlines, missed time points, and mistakenly joined cells all substantially affect how reported volumes vary with time.

Correctly segmenting asymmetrically dividing cells, such as budding yeast, is particularly challenging because the imbalance in the size of the mother and bud gives each distinct appearances and behaviours. Even when constrained in a microfluidic device, small buds imaged in a single Z section may appear to overlap with their mother and neighbouring cells (Fig. 1a & 1b). Even if cells can be semantically separated, then the area of either the bud or the neighbouring cells is often underestimated and the bud may even be undetected (Figure 1—figure supplement 1). Buds also move more in the Z-axis compared to mother cells, changing how they appear in bright-field images (Fig. 1c). Depending on the focal plane, a bud may be difficult to detect even by eye.

**Figure 1:**
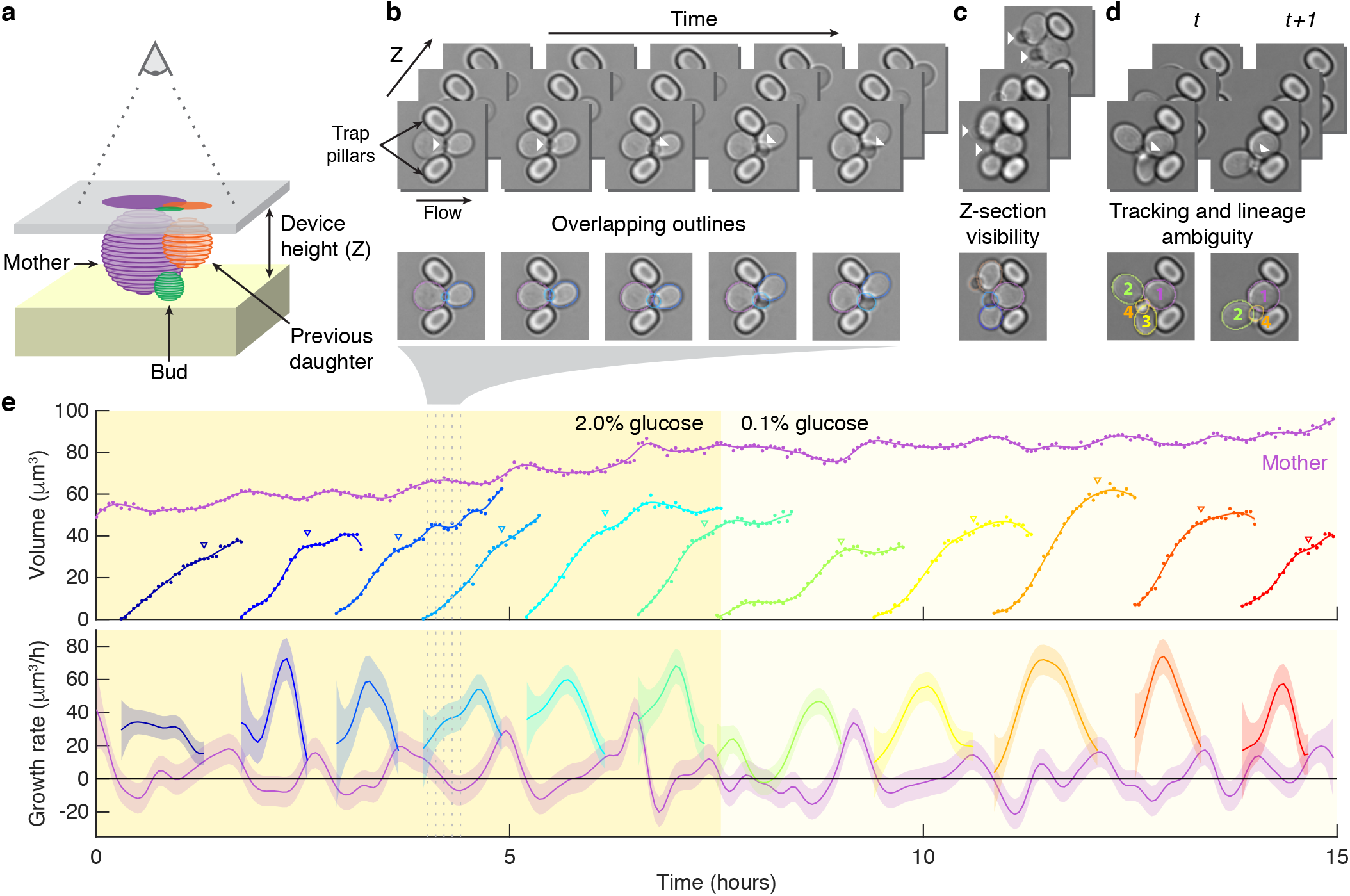
Reliably identifying individual cells makes automating label-free segmentation of budding cells challenging. **a** A schematic of a budding cell constrained in a microfluidic device showing how a mother cell can produce a bud beneath the previous daughter. The microscope, denoted by the eye, sees a projection of these cells. **b** A time series of bright-field images of budding yeast trapped in an ALCATRAS microfluidic device [38], in which a growing bud (white arrowheads) overlaps with both its sister and mother. On the duplicated images below, we show outlines produced by BABY. **c** Bright-field images of growing buds (white arrowheads) taken at different focal planes demonstrate how the appearance of small buds may change. **d** Cells can move substantially from image to image. Here medium flowing through the microfluidic device causes a cell to wash out between time points and the remaining cells to pivot. We show the correct lineage assignment with white arrowheads and the correct tracking by the numbers within the BABY outlines. **e** A time series of a mother (purple) and its buds/daughters for a switch from 2% to 0.1% glucose is shown using volumes and growth rates estimated by BABY. Bud growth rates are truncated to the predicted time of cytokinesis (triangles). Shaded areas are twice the standard deviation of the fitted Gaussian process.

Our BABY algorithm, however, has high reliability (Fig. 1e). We leverage the relative simplicity of the U-net [35], but extend this semantic approach to identify overlapping cells by combining custom training targets with a post-processing algorithm to segment instances.

To identify potentially overlapping cells from the semantic output of a convolutional neural network (CNN) like the U-net (Fig. 2a), we noted that the frequency and extent of overlap varies with cell size (Appendix 3 Fig. 2). Therefore by dividing training masks – filled cell outlines – into categories of different sizes, we could define training images for each size category that reduced whether cells overlap. We further decreased overlaps by applying and optimising a morphological erosion to the training masks (Fig. 2b; Appendix 3 Figs. 2 & 1). With these data, we trained a multi-target CNN whose output then predominantly comprised separated interiors of cells (Fig. 2c).

**Figure 2:**
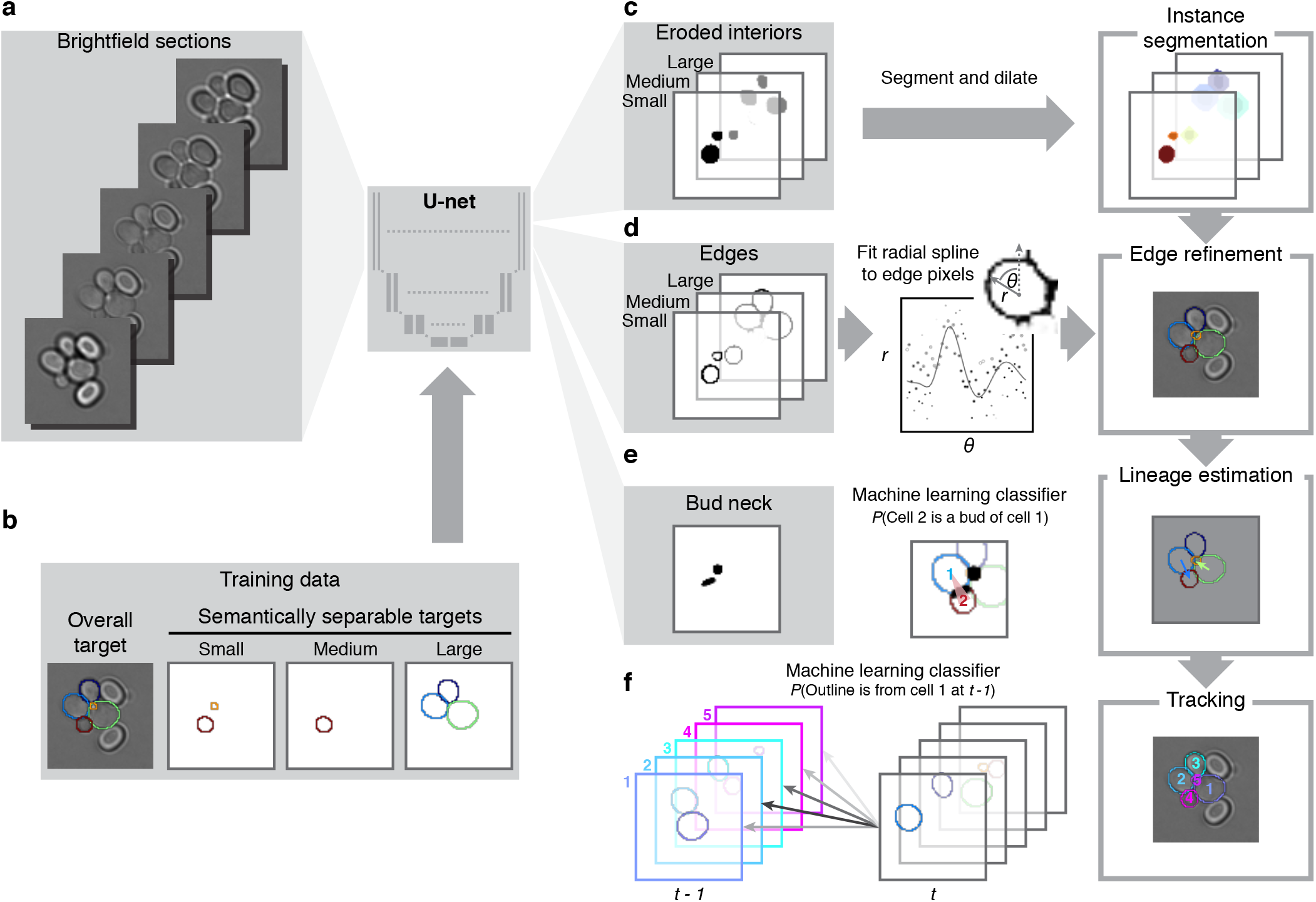
BABY uses multiple bright-field Z-sections, a multi-target convolutional neural network followed by instance segmentation, and two machine learning classifiers to identify cells and their buds reliably from image to image. **a** Multiple bright-field Z-sections are input into a multi-target U-net CNN. **b** The curated training data comprises multiple outlines that are categorised by size to reduce overlap within each category. **c** The CNN is trained to predict a morphological erosion of the target masks, which act as seeds for segmenting instances. **d** Edge targets from the CNN are used to refine each cell’s outline, parameterised as a radial spline. **e** A bud-neck target from the CNN and metrics characterising the cells’ morphologies are used to estimate the probability that a pair of cells is a mother and bud with a machine-learning classifier. **f** Another classifier uses the same morphological metrics to estimate the probability that an outline in the previous time point matches the current one.

Using a four-layer U-net, we achieved high accuracies for predicting these targets early in training and with relatively few training images (588 trap images – 1813 annotated cells in total; Appendix 3 Fig. 3). Using standard image processing, we then readily identified cell instances and applied a morphological dilation appropriate for each size category to recover the training masks (Appendix 3 Fig. 4).

To alleviate any loss in resolution from the morphological erosion used to generate the training masks, we incorporated the training masks’ edge pixels as an additional set of training targets for each size category (Fig. 2d). Like the StarDist [37] and DISCO algorithms [30], we parameterise masks using a radial representation: it naturally describes the elliptical shape of a yeast cell but also specifies any star-convex shape, including non-convex pinched ones. Specifically, we fit radial splines with 4-8 rays depending on the cell’s size to a re-weighted version of the edge pixels and use outlines estimated from the CNN targets of the cells’ interiors as initial guesses (Appendix 3 Fig. 5). The resulting masks improve the intersection-over-union score on test images (Figure 3—figure supplement 1). Importantly, our method successfully detects and segments buds that overlap with adjoining mother and daughter cells (Fig. 1b & 1c).

Other features further improve performance. We developed a powerful graphical user interface (GUI) to label and annotate overlapping cells with radial splines (Appendix 2 Fig. 1). The GUI allows concurrent viewing of the same cell across both Z-sections and time and so enables errors in segmentation and tracking errors to be rapidly identified and fixed. We therefore focused on challenging cases, iteratively cycling between training and curating in what was effectively active learning. For training, we developed scripts that optimise the hyper-parameters, and so no specialist knowledge is required to change to other imaging modalities and cell types. Finally, although BABY can predict overlapping outlines from a single 2D image, performance is improved by using multiple Z-sections (Figure 3—figure supplement 1).

The algorithm is highly accurate (Fig. 3a). For larger cell sizes, BABY performs comparably with both our previous algorithm DISCO [30], which segments instances, and a recent algorithm YeaZ [34], which uses a U-net. For smaller cell sizes, there is substantial improvement. By using size categories, BABY overcomes the difficulties from the imbalance in size of mothers and buds and identifies buds overlapping with mother cells that both DISCO and YeaZ therefore miss.

### Using machine learning to track lineages robustly

To determine growth rates accurately, estimates of both the volumes of the mother and the bud are necessary because the majority of growth occurs in the bud [13, 39]. Not only must we track cells from one time point to the next, but also correctly identify, track, and assign buds to their mothers (Appendix 4 Fig. 4).

Budding raises multiple issues. With their small initial size, buds frequently first appear surrounded by cells and displace their neighbours as they grow (Fig. 1b), potentially making assigning a bud to its mother ambiguous (Fig. 1d). Additional challenges arise when the connected mother and bud react to the flow of medium. Buds may pivot around the mother, and such movements are especially prevalent, and can even involve multiple cells (Fig. 1d), in microfluidic devices that allow long-term imaging by washing away daughter cells. If tracked incorrectly, a pivoting bud is likely to be treated as newly detected – both overestimating budding rates and underestimating lag times.

To help assign buds to mothers, we included an additional ‘bud-neck’ training target for the CNN (Fig. 2e; Appendix 4) because cytokinesis is sometimes visible as a darkening of the bud neck in bright-field images. We then estimate the probability that a pair of cell outlines is a mother and bud using a machine-learning classifier trained on both morphological features derived from the segmented outlines and the CNN’s predictions of bud-necks (Appendix 4 Fig. 3). Although the CNN’s performance in this task is lower than it is for identifying cells (Appendix 3 Fig. 3), the mother-bud classifier achieves a precision of 80% on test data.

We construct the most probable lineage tree by first identifying cells that are likely to be buds and then assigning their mothers. For each bud, we pick its mother cell as the cell that has the highest accumulated probability of being in a pair with the bud, where the probability is accumulated over all previous images containing the bud and the potential mother cell (Appendix 4).

We use another classifier to track cells between time points. This classifier is trained on the morphological properties of the cell outlines with only weak constraints on distance – the cells should only be near a microfluidic trap – allowing pivots (Fig. 1d) to be correctly tracked. Other tracking algorithms, in contrast, typically rely on distance, either explicitly by using the Hungarian algorithm to assign pairs of cells between images [25, 28, 34] or implicitly by seeding the detection of tracked cells either forwards [30] or backwards [31] in time. Using the classifier we find the probability of a match for all pairs of outlines between two time points (Fig. 2f; Appendix 4) and identify unmatched outlines too – those that have no pairings above a threshold probability. Such outlines are treated either as new buds or as disappearing or appearing cells because cells may be washed into or out of the field of view by the medium. To be resilient to transient errors in segmentation and in acquisition, like a loss of focus, we aggregate tracking predictions over the last three time points and defer to a classifier with stronger constraints on distance if assignment is ambiguous (Appendix 4).

Comparing with the DISCO [30] and YeaZ [34] algorithms, BABY finds more than three times as many complete or near-complete tracks (Fig. 3b). DISCO uses previously detected cells as priors for tracked cells in subsequent time points, and YeaZ uses the Hungarian algorithm to assign pairs that minimise differences in space and morphology. Each algorithm was assessed against manually curated data by calculating the intersection-over-union score (IoU) between cell masks in a reference track with those in a predicted track. We report the track IoU – the time points where the masks match relative to the total duration of both tracks. Where multiple predicted tracks could match a reference track, we use the match with the highest track IoU. Any predicted tracks left unassigned receive a track IoU of zero. BABY excels because it detects buds earlier, which both increases the track IoU and prevents buds from being tracked to daughters rather than mothers.

We are unaware of other algorithms that assign lineages from bright-field images of budding yeast and so compared estimates of doubling times derived by BABY with those from manual curation (Fig. 3c). The mother’s time between budding events, *T*_M_, is expected to be shorter than the time a newly born daughter takes to produce its first bud, *T*_D_ [39], and so we consider both types of doubling separately (Fig. 3c). There are fewer results for *T*_D_ because observing a daughter cell that buds before being lost in the flow is rare – the microfluidic device is designed to retain mothers not daughters [38]. Although the estimates of *T*_D_ are consistent, BABY does not give estimates for all curated daughters, usually because some buds forming on daughter cells appear for only a few time points before both daughter and bud are washed away.

**Figure 3:**
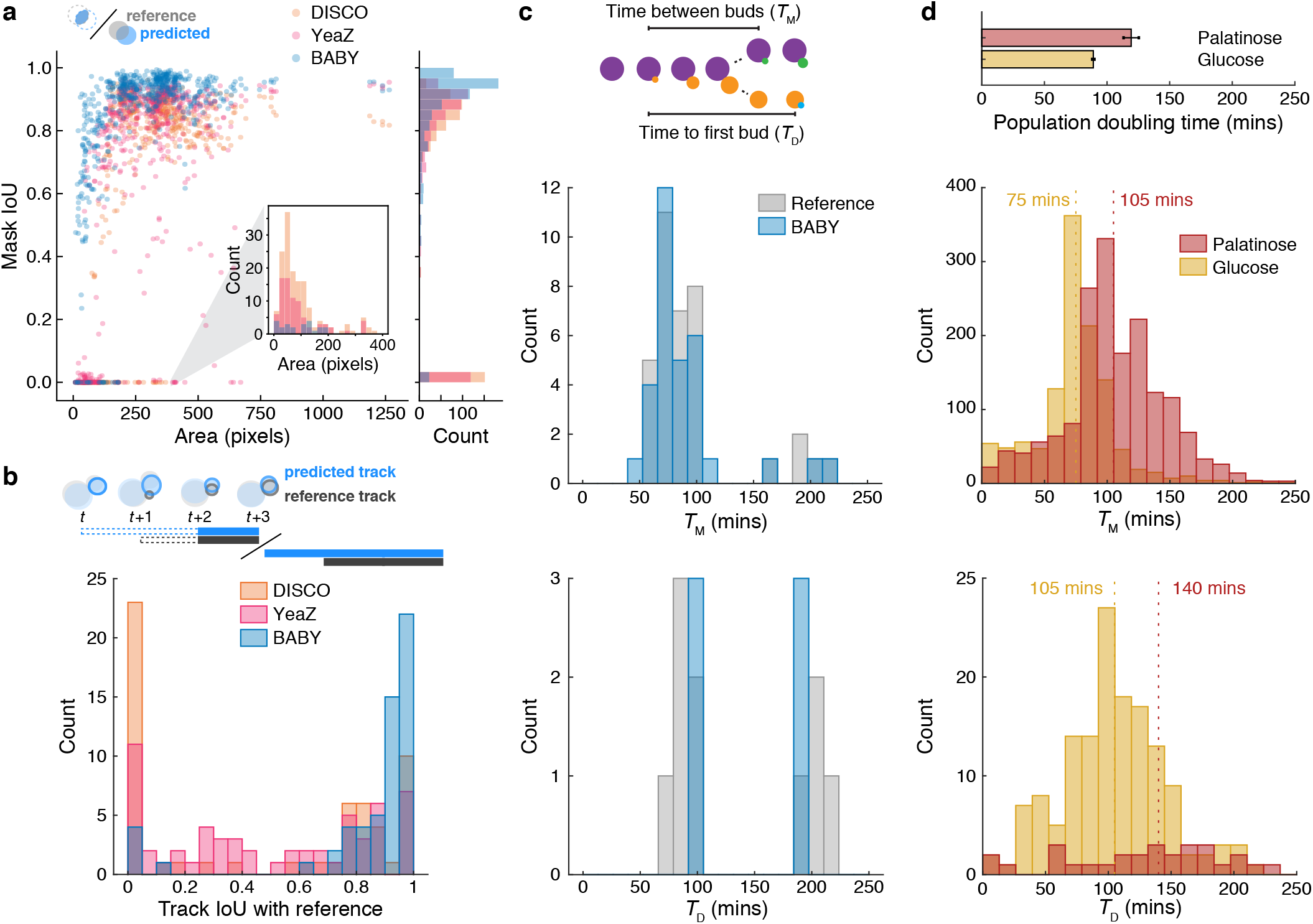
BABY outperforms existing algorithms for segmenting and tracking as well as accurately estimating doubling times. **a** Comparing the intersection-over-union (IoU) score between manually curated masks of single-cell outlines and masks predicted by the BABY, YeaZ and DISCO algorithms shows that BABY performs best, particularly for buds and small cells. **b** BABY tracks best when comparing the distribution of track IoUs between manually curated single-cell tracks and predicted tracks. **c** BABY accurately predicts both the times between budding events, *T*_M_, and the times from birth to a cell’s first budding event, *T*_D_. **d** The predicted distributions of *T*_M_ and *T*_D_ shift to longer times, as expected, when steady-state growth in 2% glucose is compared to 2% palatinose, a poorer carbon source. We estimate too the population doubling times [39] using the median times (upper bar plot with mean and 95% confidence interval from 10 bootstraps of the doubling time distributions).

As an additional test, we ran BABY for images of cells growing in palatinose and in glucose (Fig. 3d). Estimates of *T*_M_ are on average shorter than *T*_D_, and, at least for glucose where we have more division events from the faster growth, we observe that the distribution of *T*_D_ is broader than that for *T*_M_.

A useful control is to compare the population’s doubling time with that in batch growth. We estimated the doubling time from the medians of *T*_M_ and *T*_D_ with the formula of Hartwell and Unger [39]. The results for glucose (Fig. 3d) agree with our measurements of batch growth (Figure 3—figure supplement 2). BABY, however, estimates a shorter doubling time in palatinose, a sugar similar to maltose. In batch growth, the concentration of the carbon source is not constant like it is in a microfluidic device, and cells both prepare for and switch from fermentation to respiration [40], slowing growth. This switch complicates comparing with constant conditions and is likely to happen for higher concentrations of palatinose than glucose because cells strongly prefer to ferment glucose.

### Estimating growth rates

From the time series of segmented cells, we estimate instantaneous single-cell growth rates as time derivatives of volumes. We use a conical method to find each cell’s volume from its outline in an image [27] (Fig. 1e) and then a Gaussian process to both smooth the time series of volumes and to estimate its time derivative [41].

We independently estimate the growth rates of mothers and their buds. Both are informative. Like others [13, 42], we observe periodic changes in growth rate across the cell cycle (Fig. 1e).

### BABY predicts cytokinesis from bright-field images

To better characterise how growth rate varies during the cell cycle, we ran experiments for cells with the gene MYO1 tagged with Green Fluorescent Protein. As part of the actin-myosin contractile ring, Myo1 localises to the bud neck in late G1 and delocalises at cytokinesis [43] (Fig. 4a), marking the birth of a daughter. We could therefore identify the G1 and S/G2/M phases of the mother cells and the first G1 of independent daughters (Appendix 5 Fig. 1). By aligning the growth rates at cytokinesis, we observed two phases of growth (Fig. 4b). The bud dominates growth during S/G2/M, with its growth rate peaking approximately midway through (Fig. 4b) [13]. We further observe that the bud’s average growth rate positively correlates with the volume of the new daughter it becomes (Figure 4—figure supplement 1). The mother’s growth rate, in contrast, peaks during G1.

**Figure 4:**
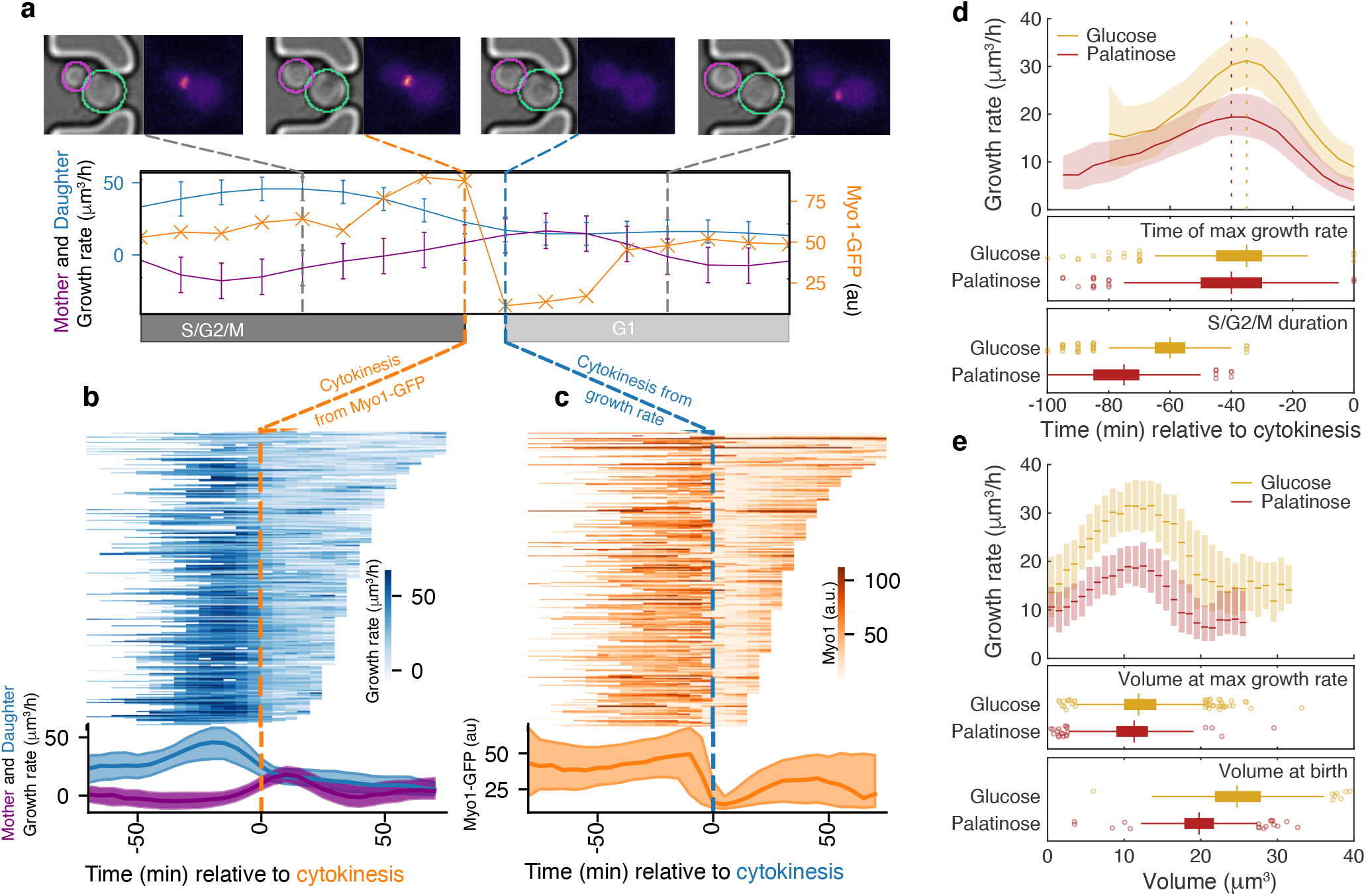
A peak in growth rate of the bud is predictive of cytokinesis and implicated in size control. **a** Using MYO1-GFP as a marker, we split individual cell cycles of mother cells into G1 and S/G2/M phases. We show bright-field and fluorescence images for a representative cell during one cell cycle. Myo1’s fluorescence at the bud neck abruptly disappears at cytokinesis. The cell’s growth rate (purple) and that of its bud (blue) are found from their volumes and shown with a measure of Myo1’s localisation (orange). We estimate the point of cytokinesis from the drop in Myo1 fluorescence (orange dashed line) and also from the mother’s and bud’s growth rates (blue dashed line). **b** The time series with shading indicating growth rates for the buds/daughters of 268 cells growing initially in 2% and switched to 0.1% glucose are shown as horizontal lines. Cells are aligned by the time of the drop in Myo1’s fluorescence. The corresponding median growth rates of mother and bud/daughter are plotted below, with the interquartile range shaded. The longer cell cycles likely occur during the glucose switch. **c** We show the fluorescence signal for the same cells, but align by the point of cytokinesis predicted from the mother’s and daughter’s growth rates. For almost all cells, Myo1’s fluorescence drops near the predicted time of cytokinesis. **d** Although buds grow faster in richer media, the time of the maximal growth rate relative to cytokinesis is approximately constant, unlike the durations of the mothers’ S/G2/M phases. Cells are grown in either 2% glucose or 2% palatinose. We show median growth rates with the interquartile range shaded. **e** Binning median growth rates according to volume, with the interquartile range shaded, shows that the buds’ volumes when their growth rate is maximal are similar in both carbon sources, although those at birth are not.

We determined that we could predict cytokinesis from growth rate because at cytokinesis the mother’s and bud’s growth rates often reach a similar magnitude (Appendix 5). The predictions coincide with the drop in Myo1’s fluorescence as expected (Fig. 4c) and hold in different media, with the Myo1-based and predicted points of cytokinesis having a Pearson correlation coefficient above 0.94 (Appendix 5 Fig. 2).

### Nutrient modulation of birth size occurs after the peak in growth rate

This tight coordination between growth rate and cytokinesis suggested that the peak in growth rate preceding cytokinesis marks a regulatory transition. Comparing growth rates over S/G2/M for buds in a rich versus a poor carbon source, we found that the maximal growth rate occurs at similar times for both carbon sources despite substantial differences in the durations of the S/G2/M phases (Fig. 4d).

Daughters born in rich media are larger than those born in poor media, and some of this regulation is known to occur during S/G2/M [5, 6, 24]. Understanding the mechanism, however, is confounded by the longer S/G2/M phases in poorer media (Fig. 4d) [24], which counterintuitively gives daughters that should be smaller longer to grow.

Given that the time between maximal growth and cytokinesis appears approximately constant in different carbon sources (Fig. 4d), we hypothesised that the growth rate falls because the bud has reached a critical size and that size is regulated during the subsequent period of falling growth to reach the desired daughter size. The faster growth in richer carbon sources would generate larger daughters. Consistently, the buds’ growth rates peak when the buds reach a similar size in both carbon sources, whereas their sizes at birth are distinct (Fig. 4e). Size regulation in S/G2/M is therefore likely to be implemented as cells approach cytokinesis.

### Changes in ribosome biogenesis precede changes in growth

An important advantage of the BABY algorithm is that single-cell growth rates can be estimated without fluorescence markers, freeing fluorescence channels for other reporters. Here we focus on Sfp1, a transcription factor that helps coordinate ribosome synthesis with the availability of nutrients [6].

Sfp1 promotes the synthesis of ribosomes by activating the ribosomal protein (RP) and ribosome biogenesis (RiBi) genes [6, 44] and enters the nucleus in response to two conserved nutrient-sensing kinases – TORC1 and PKA [6, 45, 46] (Fig. 5a). In steady-state conditions, levels of ribosomes positively correlate with growth rate [47], and we therefore assessed whether Sfp1’s activity predicts changes in instantaneous single-cell growth rates.

**Figure 5:**
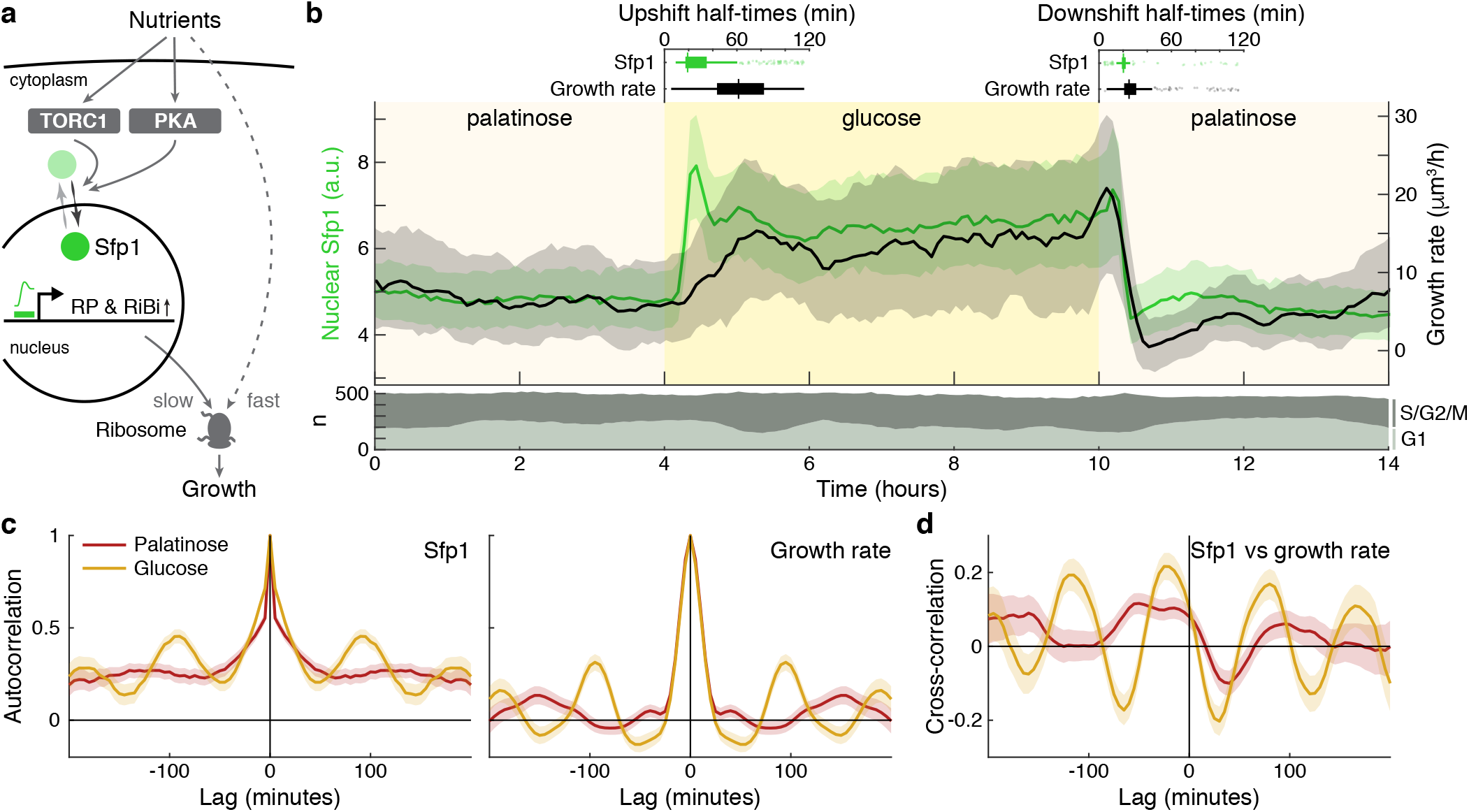
The dynamics of the ribosomal regulator Sfp1 anticipate changes in single-cell growth rates. **a** The transcription factor Sfp1 is activated by the kinases TORC1 and PKA when extracellular nutrients increase and promotes synthesis of ribosomes and so higher growth rates. **b** Growth rate follows changes in Sfp1’s nuclear localisation if nutrients decrease but lags if nutrients increase. We show the median time series of nuclear Sfp1-GFP in mothers (green) and the summed bud and mother growth rates (black) for cells switched from 2% palatinose to 2% glucose and back. The interquartile range is shaded. Data was filtered to those cell cycles that could be unambiguously split into G1 and S/G2/M phases by a nuclear marker, and we display the number in each phase in the area plot. Above the switches, we show box plots for the distributions of half-times: the time of crossing midway between the minimal and maximal values. **c** The mean single-cell autocorrelation of nuclear Sfp1 and the summed mother and bud growth rates are periodic because both vary during the cell cycle. We calculate the autocorrelations for constant medium using data four hours before each switch (Appendix 6). The 95% confidence interval is shaded. **d** The mean cross-correlation between nuclear Sfp1 and the summed mother and bud growth rate shows that fluctuations in Sfp1 precede those in growth.

Shifting cells from glucose to the poorer carbon source palatinose and back again, we observed that Sfp1 responds quickly to both the upshift and downshift and that growth rate responds as quickly to downshifts but more slowly to upshifts (Fig. 5b). As a target of TORC1 and PKA, Sfp1 acts as a fast read-out of the cell sensing a change in nutrients [20]. In contrast, synthesising more ribosomes is likely to be slower and explains the lag in growth rate after the upshift. The fast drop in growth rate in downshifts is more consistent, however, with ribosomes being deactivated, rather than their numbers being regulated. Measuring the half-times of these responses (Fig. 5b boxplots), there is a mean delay of 30 2 minutes (95% confidence; *n* = 245) from Sfp1 localising in the nucleus to the rise in growth rate in the upshift. This delay is only 8 1 minutes (95% confidence; *n* = 336) in the downshift, and downshift half-times are less variable than those in upshifts, consistent with fast post-translational regulation. Although changes in Sfp1 consistently precede those in growth rate, the higher variability in half-times for the growth rate is not explained by Sfp1’s half-time (Pearson correlation 0.03, *p* = 0.6).

By enabling both single-cell fluorescence and growth rates to be measured, BABY permits correlation analyses [21] (Appendix 6). Both Sfp1’s activity and the growth rate vary during the cell cycle. The autocorrelation functions for nuclear Sfp1 and for the growth rate are periodic with periods consistent with cell-division times (Fig. 5c): around 90 minutes in glucose and 140 minutes in palatinose for Sfp1 and 95 minutes and 150 minutes for the growth rate. If Sfp1 acts upstream of growth rate, then its fluctuations in nuclear localisation should precede fluctuations in growth rate. Cross-correlating nuclear Sfp1 with growth rate shows that fluctuations in Sfp1 do lead those in growth rate, by an average of 25 minutes in glucose and by 50 minutes in palatinose (Fig. 5d). Nevertheless, the weak strength of this correlation suggests substantial control besides Sfp1.

During the downshift, we note that the growth rate transiently drops to zero (Fig. 5b), irrespective of a cell’s stage in the cell cycle (Figure 5—figure supplement 1), and there is a coincident rise in the fraction of cells in G1 (Fig. 5b bottom), suggesting that cells arrest in that phase.

### Using growth rate for real-time control

With BABY, growth rate can be used as a control variable in real time because BABY’s speed and accuracy enables cells to be segmented and their growth rates estimated during an experiment (Fig. 6a). As an example, we switched the medium to a poorer carbon source and used BABY to determine how long cells should be kept in this medium if we want approximately 50% the cells to have resumed dividing before switching back to the richer medium (Appendix 7). After five hours in glucose, we switched the carbon source to ethanol (or galactose – Figure 6—figure supplement 1). There is a lag in growth as cells adapt. Using BABY, we automatically determined the fraction of cells that have escaped the lag at each time point – cells that have at least one daughter whose growth rate exceeds a threshold (Fig. 6b). This statistic is read by the software running the microscope, and the switch back to glucose is triggered when 50% of the cells have escaped (Fig. 6c). All cells resume dividing in glucose and initially grow synchronously because of the rapid change of media. This synchrony is most obvious in those cells that did not divide in ethanol (Fig. 6c).

**Figure 6:**
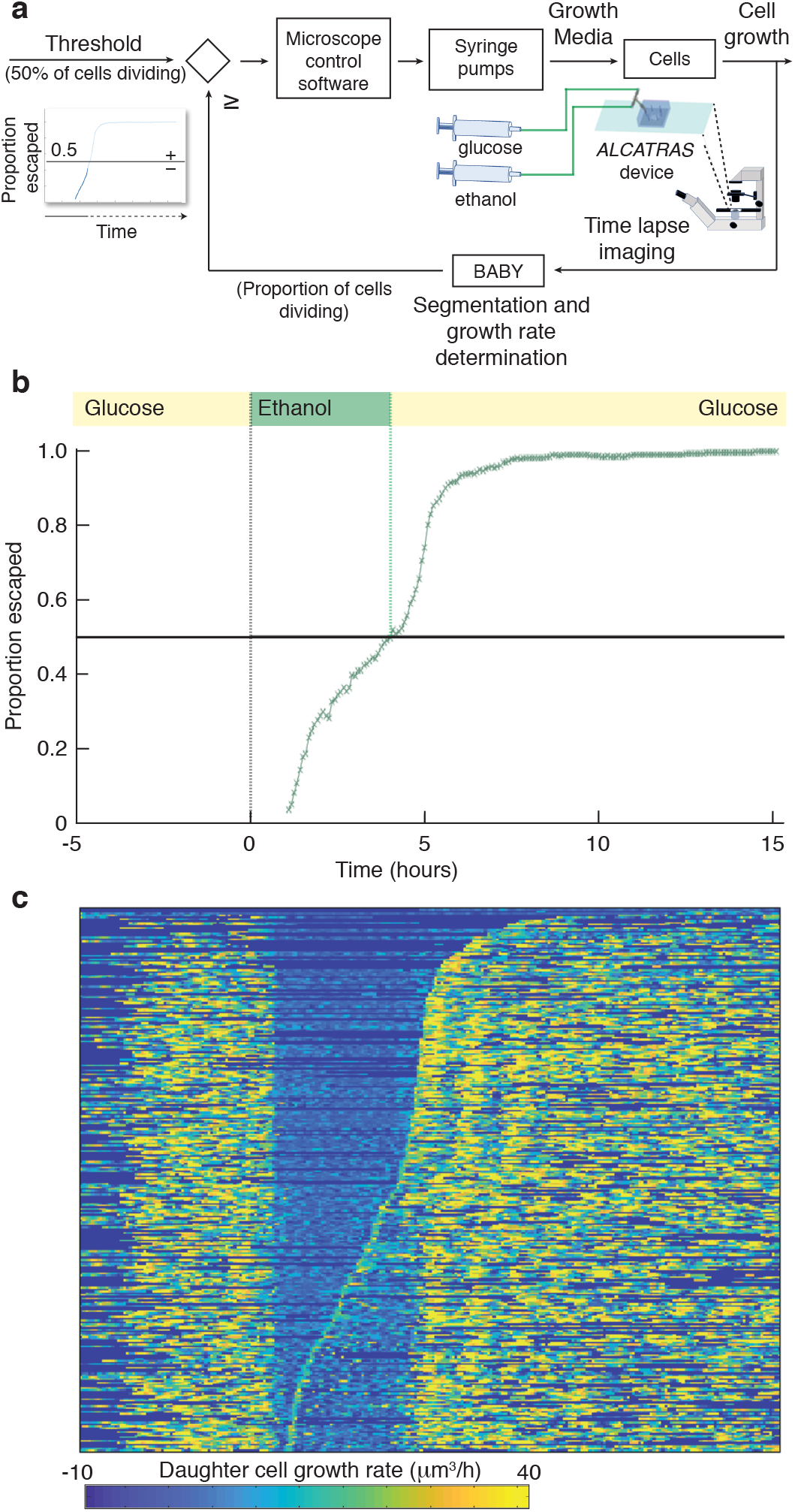
BABY allows growth rate to be used as a variable for real-time control. **a** Integrating BABY in real time with an experiment allows the cells’ growth rate to control changes in media. Cells growing in the ALCATRAS microfluidic device are fed from one of two syringe pumps containing differing media. Following five hours in 0.5% glucose, growth is arrested by a switch to 2% ethanol, a poorer carbon source. The images collected are analysed by BABY and growth rates are determined. When the majority of cells have resumed dividing, detected by the growth rate of at least one of their daughters exceeding 15 *μ*m^3^/hour, the software triggers a change in pumping and returns glucose to the device. **b** The fraction of cells that have escaped the lag and resumed dividing increases with the amount of time in ethanol. All cells divide shortly after glucose returns. **c** The growth rates of the buds for each mother cell drop substantially in ethanol and resume in glucose. Each row shows data from a single mother cell with the daughters’ growth rate indicated by the heat map. Rows are sorted by the time each cell resumes dividing in ethanol.

This proof-of-principle shows that BABY can be used in more complex feedback control, where a desired response is achieved by comparing behaviour with a computational model to predict the necessary inputs, such as changes in media [48–53]. Unlike previous approaches though, which typically measure fluorescence, BABY not only allows single-cell fluorescence but also growth rates to be control variables, and growth rate correlates strongly with fitness [1].

## Discussion

Here we presented BABY, a freely available algorithm to routinely extract growth rates from label-free images obtained by time-lapse microscopy. BABY tackles previously unaddressed challenges in analysing images of budding cells and provides fully automated tracking of lineages of budding yeast – from identifying cells to estimating births. We introduce both a segmentation algorithm that allows instances to overlap and general machine-learning methods to track and assign lineages. These approaches individually and together provide substantial gains in performance. As a result, BABY reliably estimates time series of cellular volumes and consequently instantaneous single-cell growth rates. There is no prohibitive penalty on speed, and BABY performs real-time processing.

BABY improves segmentation largely because it allows cells to overlap. Samples for microscopy are often prepared to encourage cells to grow in a monolayer [11], but growth can be more complex because cells inevitably have different sizes. We observe substantial and frequent overlaps between buds and neighbouring cells in ALCATRAS microfluidic devices [38]. Inspecting images obtained by others, we believe overlap is a common, if undeclared, problem, occurring during growth in CellASIC devices [31, 34], against an agar substrate [15, 28], in a microfluidic dissection platform [26], and by surface attachment [54]. BABY alleviates having to constrain cells to completely exclude their neighbours, at least when determining growth rates. Moreover we anticipate that BABY will segment mixtures of cells with a range of sizes even if they do not bud, such as persister cells that transition at different rates from smaller to larger morphologies [55]. For substantial growth beyond a monolayer, however, a different approach with full threedimensional imaging and segmentation is likely required [32].

Our algorithm uses a convolutional neural network that is smaller than the typical networks, with performance relying instead on the choice of output targets. U-nets usually involve five contracting layers [32, 34, 35], whereas we found four layers were optimal, presumably because our problem is of lower than usual complexity. Unlike others [30, 56], we need not explicitly ignore features in the image because the network quickly learns to disregard both the traps in ALCATRAS devices and any debris.

The neural network represents cell edges at their native resolution, unlike alternative algorithms that also allow instances to overlap: the Mask-RCNN scales regions of interest to a standard low-resolution image [36] and StarDist represents an object’s edge pixels as star-convex shapes with a constant and necessarily limited number of rays at fixed angles [37]. Though BABY’s final masks for segmentation are also star-convex, we optimise both the number of rays and their angles to fit the high-resolution edge and use smoothing splines for the cell outlines. Instances are therefore represented in an expanded morphological space, which aligns the rays with how a cell is oriented. With extra training data, we expect BABY to segment atypical morphologies, such as those of ageing cells, but for more deviant shapes specialised algorithms may be required [34].

We use a flexible, generalizable approach to tracking cells and lineages. To identify the same cell from one image to the next, tracking algorithms often use both Euclidean distance and morphological similarity, with the weighting between the two tuned to the type of image [34, 57]. We generalise this approach by applying machine learning to assign pairs. We thus both formalise how weights are determined and find a probability for each assignment. With these probabilities, we are able to distinguish rigorously between a cell that is correctly tracked and a new cell replacing another and so avoid having to arbitrarily set thresholds. Because buds appear as independent objects that need pairing with mother cells, we apply the same approach to tracking lineages and obtain high accuracies by using features derived from the neural network’s prediction of bud necks.

Once we have generated time series of cellular volumes, we make only weak assumptions about the nature of growth to estimate instantaneous growth rates. Single-cell growth rates in yeast have been variously modelled – for example, as bilinear [13, 24, 25, 42] and as exponential [7, 15, 23, 58] – but that choice has implications for size control [8]. Instead we use Gaussian processes to estimate growth rates [41] and so make assumptions only on the class of functions that describe growth rather than their precise functional form. As well as consistently estimating errors and interpolating to new time points, we show that this non-parametric analysis yields detailed instantaneous growth rates both across the cell cycle and in response to shifts in nutrients.

A well-known challenge in automating segmentation is determining when two cells become independent [57], which we answer by exploiting characteristics of yeast’s growth. Like others [13, 42], we observe that growth rate varies during the cell cycle and in particular peaks during budded growth. This peak is sufficiently regular that we are able to use it to reliably predict the time of cytokinesis.

Our results highlight the importance of single-cell growth rates as phenomenological readouts. We hypothesise that the peak in the bud’s growth rate marks a regulatory transition, triggered when the bud’s size reaches a nutrient-independent threshold. Although the bud’s size is known to be regulated during M phase [24, 25], our data suggest a sequential mechanism that matches size to growth rate, with a nutrient-independent sizer followed by a nutrient-dependent timer. This peak in bud growth may be monitored by Gin4-related kinases [59]. We also report that activation of Sfp1 – a downstream target of TOR kinase [60] – anticipates growth rate, both within a cell-cycle and after a shift in nutrients, consistent with growth being powered by translation [6].

Much of cell biology is focused on understanding how cells respond to change [10], and watching individual cells in real time as their environment alters gives great insight [11]. Together time-lapse microscopy, microfluidic technology, and fluorescent proteins allow us to control extracellular environments, impose dynamic changes, and phenotype cellular responses over time. With BABY, we add the ability – with no extra imaging costs – to measure what is often our best estimate of fitness, single-cell growth rates. The strategies used by cells in their decision making are of high interest [61, 62]. With BABY, or comparable software, we are not only able to rank the strategies that each cell adopts by their fitness, but also to investigate the strategies used to regulate fitness itself through how cells control their growth, size, and divisions.

## Methods

### Strains and media

Strains included in the curated training images were all derivatives of BY4741 [63]. Both BY4741 Myo1-GFP Whi5- mCherry and BY4741 Sfp1-GFP Nhp6A-mCherry were derived from the respective parent in the *Saccharomyces cerevisiae* GFP collection [64] by PCR-based genomic integration of mCherry-Kan^*R*^ from pBS34 (EUROSCARF) to tag either Whi5 or the chromatin-associated Nhp6A protein. All tags were validated by sequencing. Media used for propagation and growth was standard synthetic complete (SC) medium supplemented either with 2% glucose, 2% palatinose or 0.5% glucose depending on the starting condition in the microfluidics devices. Cells were grown at 30°C.

### Microscopy and microfluidics

#### Device preparation and imaging

Overnight cultures were inoculated with low cell numbers such that they would reach mid-log phase in 13-16 hours. Cells were diluted in fresh medium to OD_600_ of 0.1 and incubated an additional 3–4 hours before loading into microfluidic devices at OD_600_ of 0.3–0.4. To expose multiple strains to the same environmental conditions and to optimise data acquisition, we use multi-chamber versions of ALCATRAS [38, 65, 66], which allow for either three or five different strains to be observed in separate chambers while exposed to the same extracellular medium. The ALCATRAS chambers were pre-filled with growth medium with added 0.05% bovine serum albumin (BSA) to facilitate cell loading and reduce clumping. All microfluidics media were passed through 0.2 *μ*m filters before use.

We captured images on a Nikon Ti-E microscope using a 60, 1.4 NA oil immersion objective (Nikon), OptoLED light source (Cairn Research) and sCMOS (Prime95B) or EMCCD (Evolve) cameras (both Photometrics) controlled through custom MATLAB software using Micro-manager [67]. The microscope and syringe pumps containing media were maintained at 30°C in a custom-made incubation chamber (Okolabs).

#### Changing the extracellular environment

For experiments in which the cells experience a change of media, two syringes (BD Emerald, 10ml) mounted in syringe pumps (Aladdin NE-1002X, National Instruments) were connected via PTFE tubing (Smiths Medical) to a sterile metal T-junction. The T-junction’s output was delivered via PTFE tubing to the microfluidic device. Initially the syringe with the first medium infused at 4 *μ*L/min while the second pump was off. To remove back pressure and achieve a rapid switch, medium was infused at 150 *μ*L/min for 40s from the second pump while the first withdrew at the same rate. The second pump was then set to infuse at 4 *μ*L/min and the first switched off. This sequence was reversed to achieve a second switch in some experiments. All changes of the flow rates and directions of the pumps were controlled through Matlab software, via RS232 serial ports.

### Data and code availability

Data is available at https://doi.org/10.7488/ds/3427 and code from

https://git.ecdf.ed.ac.uk/swain-lab/baby.

### Birth Annotator for Budding Yeast (BABY) algorithm

The BABY algorithm takes either a stack of bright-field images or a single Z-section as input and coordinates multiple machine-learning models to output individual cell masks annotated for both tracking and lineage relationships.

Central to segmenting and annotating lineages is a multi-target convolutional neural network (CNN). Each target of the CNN is semantic – binary labels are assigned to each pixel. These targets are defined to ease both segmenting overlapping instances and assigning lineages in post-processing steps. The reverse task of identifying cell instances from the semantic outputs of the CNN comprises two steps: image processing to identify candidate masks and optimising a radial spline to refine mask edges. Details are given in Appendix 3.

To track cells and lineages, we use machine-learning classifiers both to link cell outlines from one time point to the next and to identify mother-daughter relationships. The classifier converts a feature vector, representing quantitatively how two cell masks are related, into probabilities for two possible classes. For cell tracking, the probability of interest is the probability that the two cells at different time points are in fact the same cell. For assigning lineages, the probability of interest is the probability that the two cells have a mother-daughter relationship. A target of the CNN dedicated to assigning lineages is aggregated over time to determine this probability. Details are given in Appendix 4.

Scripts automate the training process, including optimising hyperparameters – like the size categories and CNN architecture – and post-processing parameters. We split the training data into training, validation, and test sets [68], with these categories consistently carried through the training pipeline. The training set is used for training the CNN and the validation set for optimising hyperparameters and post-processing parameters. The test set is excluded during this optimisation and is used only to assess performance and generalisability after training.

The algorithm is implemented in Python and depends on Tensorflow [69] for the deep-learning models, Scikit-learn [70] for machine learning, and Scikit-image [71] for image processing. The code can be run either directly from Python or as an HTTP server that processes submitted requests and enables access from other languages, such as Matlab.

### Measuring growth

#### Calculating cell volumes

To estimate cell volumes, we model a 3D cell from our 2D outline. Although the neural network uses information in the *z*-direction, the resulting segmentation is 2D not 3D, partially because 3D measurements made by epifluorescence microscopy are anisotropic. In our experiments, the distance between two *z*-positions is usually approximately three times larger than the pixel size in the *x* and *y* directions.

We use a conical method [27], which is robust to various common cell shapes, to estimate cell volume from an outline. The method makes two assumptions: that the outline obtained cuts through the widest part of the cell and that the cell is ellipsoidal. We build a cone with a base shape that is the filled segmentation outline of the cell by iteratively eroding the mask of the cell and stacking these masks in the *z* dimension. We find the volume of the cone by summing the voxels in the corresponding 3D mask. Finally, we multiply this sum by four to obtain the volume of the cell, because a cone whose base is the equatorial plane of an ellipsoid will have a volume that is a quarter of the corresponding ellipsoid’s volume [27].

#### Estimating single-cell growth rates

Depending on the need for computational speed, we use one of two methods for estimating instantaneous growth rates.

For long-term, and stored, analysis, we estimate growth rates by fitting a Gaussian process with a Matern covariance function to the volume time series for each cell [41]. We set the bounds on the hyperparameters to prevent overfitting. Maximising the likelihood of the hyperparameters, we are able to obtain the mean and first and second time derivatives of the volume, as well as estimates of their errors. The volume’s first derivative is the single-cell growth rate.

During real-time processing where the growth rate may be used to control the microscope, fitting a Gaussian process is too slow. Instead we estimate growth rates from the smoothed first derivative obtained by Savitzky-Golay filtering of each cell’s volume time series. Though faster, this method is less reliable than a Gaussian process and does not estimate errors. For time series of mothers, we use a third-order polynomial with a smoothing window of seven time points; for time series of daughters, we use a fourth-order polynomial also with a smoothing window of seven time points.

We estimate growth rates separately for mothers and their buds because both are informative. We find that the summed results are qualitatively similar to previous estimates of growth rate, which fit the time series of the combined volume of the mother and its bud [13, 42].

## Acknowledgements

We thank Yu Huo for providing the data shown in Supplementary Figure 3 and gratefully acknowledge support from the Leverhulme Trust (JMJP & PSS – grant number RPG-2018-04), the BBSRC (IF, IBNC, & PSS – grant number BB/R001359/1), and the European Union’s Horizon 2020 research and innovation programme under the Marie Skl odowska Curie grant agreement no. 764591 — SynCrop (AFM & DYA).

## Appendix 1

### Quantifying localization

During each experiment, we acquired bright-field and fluorescence images at five *z*-sections spaced 0.6 micron apart. The maximum projection of these images (the maximum pixel values across all *z*-sections) was used for quantification.

For each cell, we determined its fluorescence image by subtracting the median fluorescence of the cell’s pixels and setting all non-cell pixels to zero. This median fluorescence will be determined by a cytoplasmic pixel, and we assume that it results from autofluorescence only, which requires sufficiently low numbers of fluorescent markers.

To quantify fluorescent markers in the nucleus, we noted that fluorescence in the nucleus appears in an image as a two-dimensional Gaussian distribution because of point spreading in epifluorescence microscopes. We therefore identified the most probable location of the nucleus for each cell by convolving a Gaussian filter with the fluorescence image. The maximal value in the resulting filtered image marks the location that most closely matches this filter.

Using data from nuclei segmented via Nhp6A-mCherry reporters [20], we observed that the area of the nucleus *A*_nuc_ scales as a fraction of cell area *A*_cell_ with a scaling factor *f*_nuc_ ≃ 0.085. We used this result to estimate a standard deviation *σ* for the Gaussian filter. If the nucleus is approximately circular then its radius can be estimated as

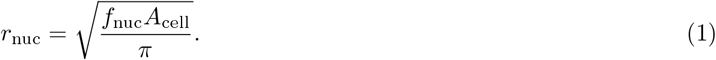

Assuming that 95% of the fluorescence in the nucleus is contained within the segmented area of nucleus, then we choose the *σ* of the Gaussian filter so that 95% of its probability is obtained by integrating over a circle of radius *r*_nuc_. Writing α = 0.95, we have

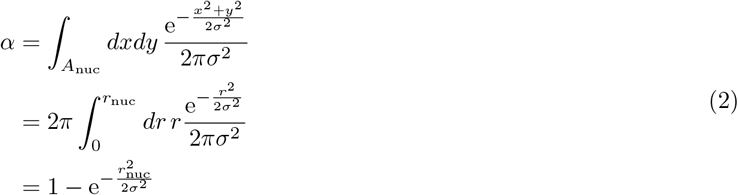

switching to polar coordinates. Using that the cumulative distribution function of the *x*^2^ distribution with two degrees of freedom is 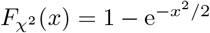, we can rearrange Eq. 2 and combine with Eq. 1 to give

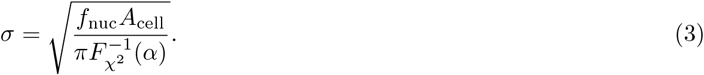

We next assume an ideal fluorescence image of the nucleus can be described by the same Gaussian filter but rescaled by some amplitude α_*f*_. If we apply the Gaussian convolution *G* to the pixel in this ideal image with maximal fluorescence, we obtain

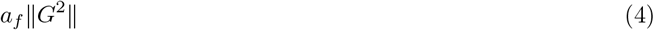

where ∥*G*^2^∥ is the sum of the squared values of the Gaussian filter. This quantity should in principle be equal to αmax(*C*), where *C* is the Gaussian filtered fluorescence image of the actual cell. Therefore

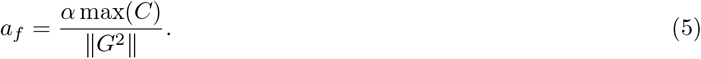

Finally, α_*f*_ is our prediction of the total nuclear fluorescence, but the concentration is more biologically relevant and, if denoted *N*, is

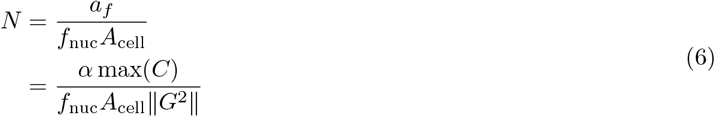

which is the measure we use.

For quantifying the localization of Myo1-GFP to the bud neck, we note that *N* is a sensitive proxy for localization and assume that it applies equally well in this case.

## Appendix 2

### Training data and a graphical user interface for curating

**Figure 1:**
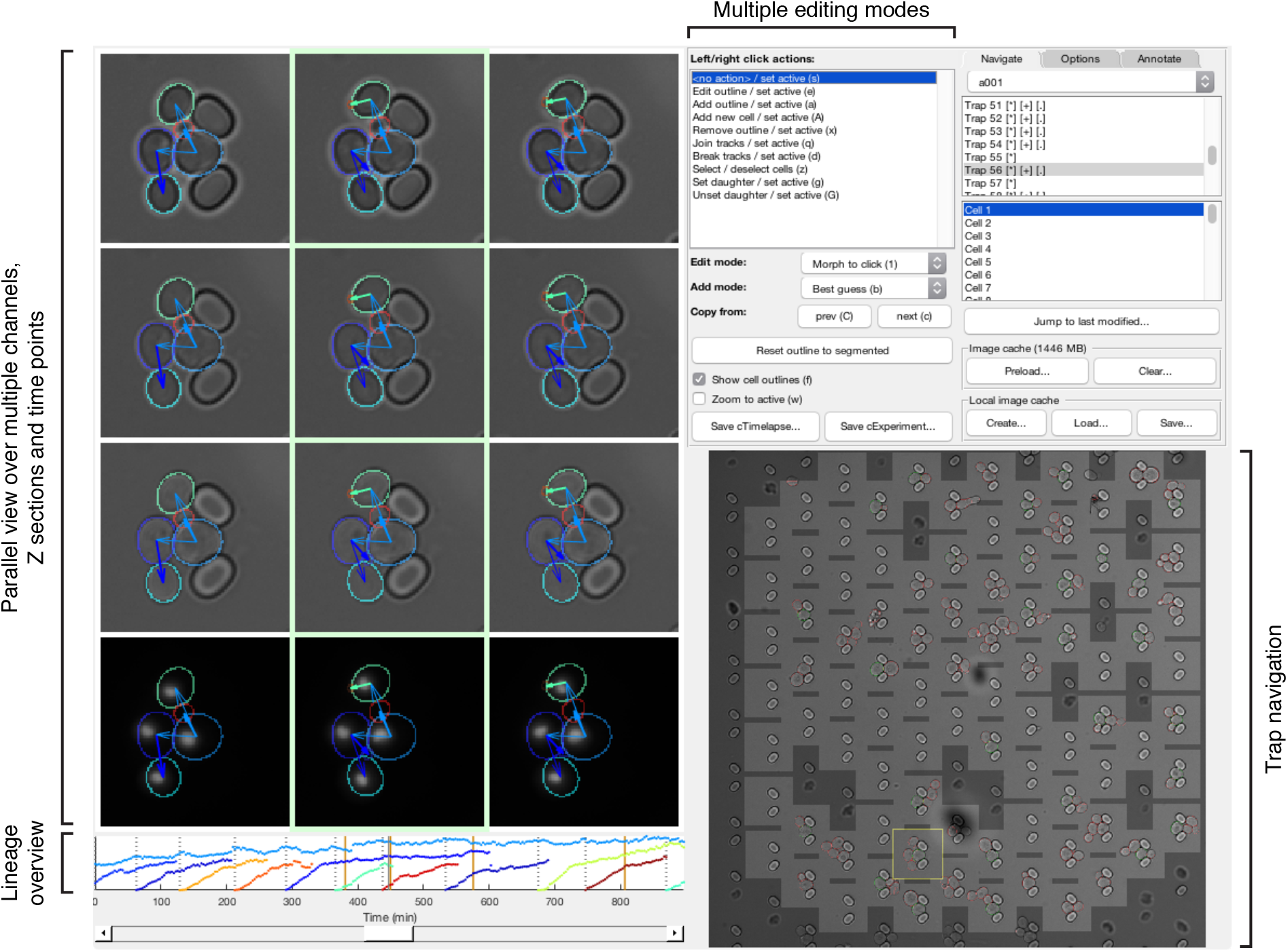
Main features of the graphical user interface used for annotation. A custom graphical user interface (GUI) was developed in Matlab for efficient annotation of overlapping cell instances, tracks and lineages over long time courses. The screen-shot shows the GUI in its horizontal layout with 3 bright-field sections and a fluorescence channel selected for parallel view. Annotated outlines and arrows indicating lineage relationships have each been toggled on for display. Up to 9 time points can be displayed in parallel; the slider at the bottom allows fast scrolling through the entire time-lapse. A time-course summary panel is displayed above the slider and has been set to show the outline areas for a mother and all its daughters. An overview image of the entire position allows navigation between traps. Multiple editing modes can be selected for manipulating annotations in the parallel view region, including modes for draggable outline editing, track merging/splitting and lineage reassignments.

Training data for the machine learning models consisted of bright-field time-lapse images of yeast cells trapped in ALCATRAS devices [38] and manually curated annotations: a bit-mask outline for each cell and its associated tracking label and lineage assignment, if any. All images were taken with five Z sections and 60 lens, including examples taken using cameras with different pixel sizes (0.182 *μ*m and 0.263 *μ*m).

To ease annotating overlapping instances, cell tracks and lineage relationships, we developed a Graphical User Interface (GUI) in Matlab that allows parallel viewing of multiple Z sections and time points (Appendix 2 Fig. 1). This parallel view helps curate buds obscured by a lack of focus and those that might be missed without simultaneously observing multiple time points. Manipulations made to outlines and tracks are mirrored to all views in real time. The interface is highly customisable, with multiple layouts available and the ability to select which sections and channels are displayed. To edit outlines for smaller cells, the level of zoom can be adjusted. Further, starting outlines can be copied across time points and interpolated forwards or backwards in time (interpolated outlines are annotated as such until they are manually adjusted).

Annotations are saved in a custom format for computational efficiency, but various export options are available. For training we exported annotations in PNG format with one image per time point. Because outlines can potentially overlap, they are tiled, with one cell instance per tile. The metadata of the PNG file is used to store track and lineage annotations.

The GUI also includes features to efficiently detect and correct rare errors. A track display panel provides visual aids to summarise tracks across the entire time course. In particular, the ‘Display mother and daughter areas’ mode uses this panel to plot the area of the currently selected cell and all of its daughters over the time course. Using this mode, many segmentation and tracking errors are highlighted as unexpected jumps in area or changes in track label (denoted in colour). We use a slider to navigate to these errors where they can be either corrected in place or saved for future curation.

Although the GUI can be used on whole images, it includes features to navigate and annotate images that can be partitioned into regions, such as the traps of our ALCATRAS devices. Then the trap navigation image shows trap locations and can be used to move between traps.

## Appendix 3

### The BABY algorithm: identifying cells and buds

#### Mapping cell instances to a semantic representation

For epifluorescence microscopy, samples are typically prepared to constrain cells in a monolayer. If the cells have similar sizes and match the height of this constraint, they will be physically prevented from overlapping. If cells are of different sizes, however, then a small cell can potentially fit in gaps and overlap with others. This phenomenon is especially prevalent for cells that divide asymmetrically, where a small daughter grows out of a larger mother.

Few segmentation algorithms identify instances of overlapping cells. Most, including recent methods for budding yeast [31, 34, 56], make the assumption that cells can be labelled semantically – each pixel of the image is identified with at most one cell. Similarly, most tools for annotating also label semantically, and consequently curated training data does not allow for overlaps [34], even when the segmentation algorithm could [33]. Our laboratory’s previous segmentation algorithm included limited overlap between neighbouring cells [30], but not the substantial overlap seen between the smaller buds and their neighbours.

**Figure 1:**
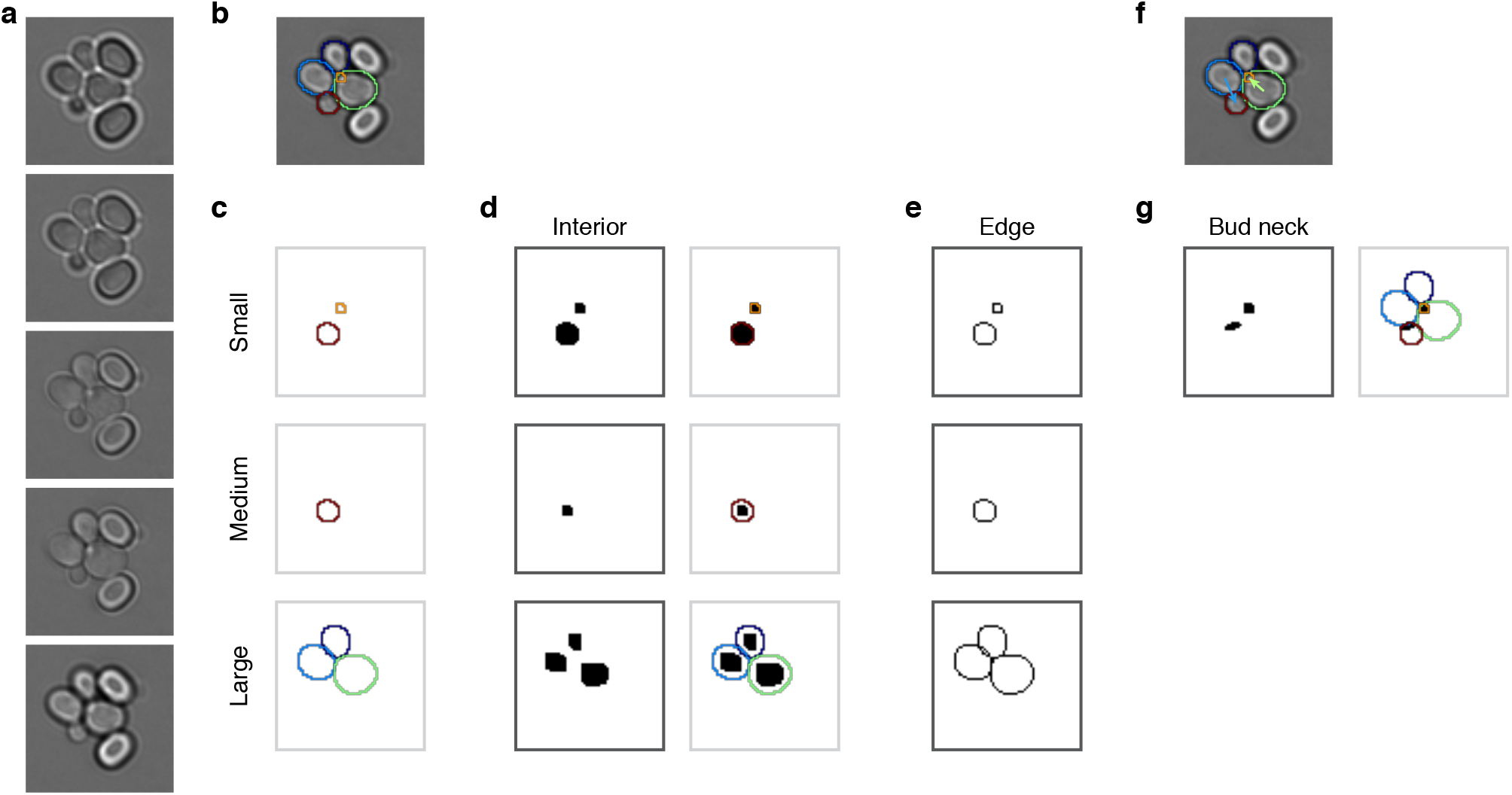
Mapping cell instances to semantic targets of a CNN. **a** Bright-field Z-sections of cells trapped in an ALCATRAS device. **b** Curated cell outlines overlaid on one bright-field section. **c** Outlines are separated into categories by size. Each category is chosen to retain some overlap with neighbouring categories. Here the red outline in the medium category is duplicated in the small category. **d** Interior targets are cell masks after different rounds of morphological erosions appropriate for each size category: no erosion for small cells, four iterations for medium, and five for large. On the right, outlines are overlaid on the target masks for reference. **e** Edge targets are the outlines for each size category. **f** The curated cell outlines of **b**, but with arrows to show the lineages assigned during curation. **g** Using these curated lineages, the ‘bud neck’ target is defined as the overlap of the bud mask with a morphological dilation of the mother mask (right).

#### Separating cells by size to disjoin overlapping cells

We rely on two consequences of the height constraint to segment overlapping instances. First, cells of different sizes show different patterns of overlap; second, cells rarely have coincident centres. Very occasionally, we do observe small buds stacked directly, one on top of the other, but neglecting these rare cases does not degrade performance. We therefore use morphological erosions to obtain semantic images by shrinking cell masks within a size category and morphological dilations to approximate the original cell outlines from each resulting connected region.

To separate overlapping cells, we define three size categories and treat instances in each category differently. Appendix 3 Fig. 1 illustrates our approach, where we segment a bud (orange outline) that overlaps a mother cell (green outline). The bud is only visible in the third and fourth Z sections of the bright-field images (Appendix 3 Fig. 1a). If used for training, the manually curated outlines in this example (Appendix 3 Fig. 1b) would be split into different size categories (Appendix 3 Fig. 1c). The bud is assigned to the small category. When the outlines in this category are filled and the image converted into a binary one (Appendix 3 Fig. 1d), the cell masks are distinct from each other. For the large category, however, the masks are not separable when immediately converted, but become so when the filled outlines are first morphologically eroded (Appendix 3 Fig. 1d). The largest size category tolerates more erosions than smaller ones, for which the mask may disappear or lose its shape.

#### Determining the size categories

Using the training data – curated masks for each cell present at each trap at each time point, we identify the size categories that best separate overlapping cells. To begin, we calculate the overlap fraction – the intersection over union – between all pairs of cell masks. Its distribution reveals that the most substantial overlaps occur between cells of different sizes (Appendix 3 Fig. 2a – upper triangle).

**Figure 2:**
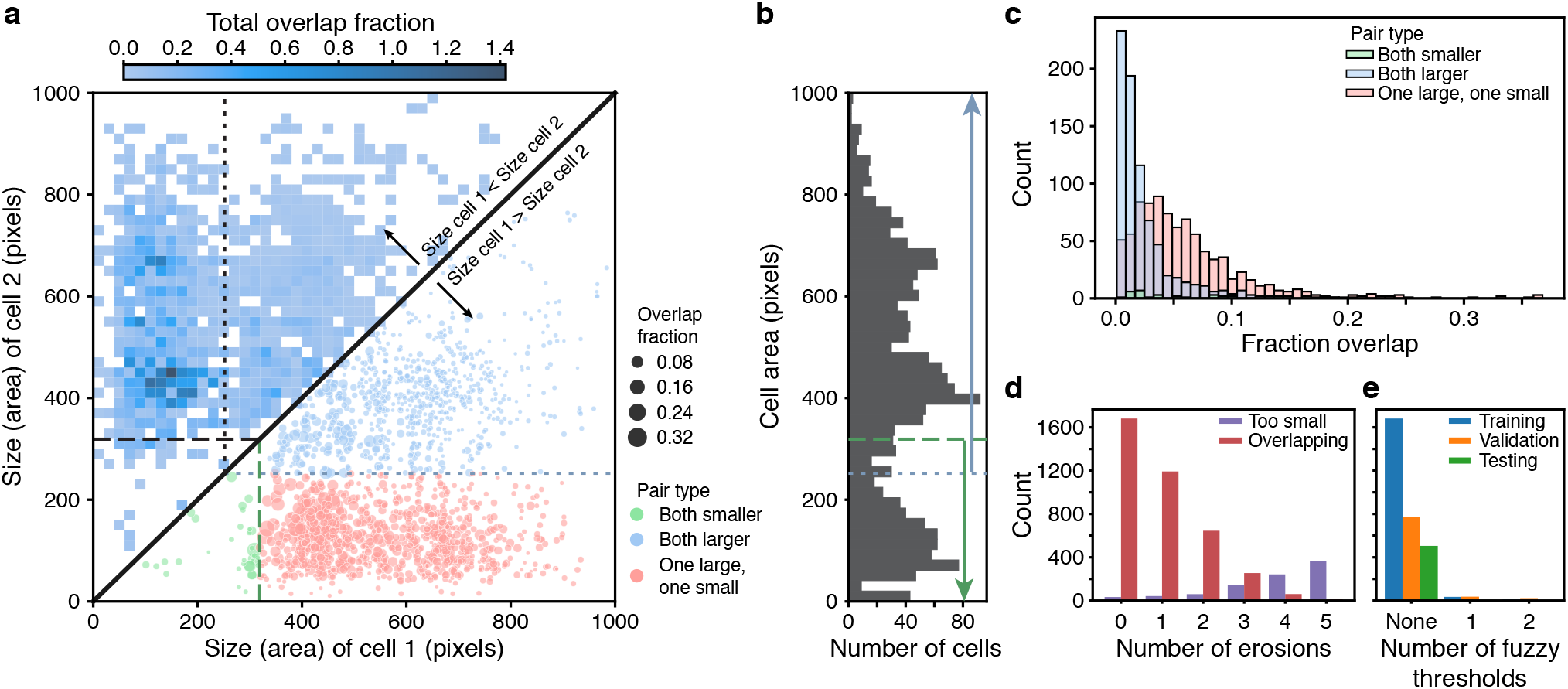
Overlaps between cells are reduced through categorising cells by size. **a** Upper triangle: plotting the overlap fraction for each pair of cells – the intersection over union of their bit masks, shows that the majority of overlaps occur for cells of different sizes. Almost all overlaps have the size of cell 2 greater than the size of cell 1 and lie off the diagonal. Lower triangle: With a single fuzzy size threshold, cells in the small category have sizes less than the upper threshold (dashed line), and cells in the larger category have sizes greater than the lower threshold (dotted line). Within each category, the overlap fractions are mostly small; between the two categories, overlaps are mostly large (excluded cells in red). By converting the bit masks into two binary images, one for each size category, rather than a single binary image, we therefore eliminate, or exclude, most of the substantial overlaps. **b** The distribution of all mask areas in the same training data for comparison. Size thresholds are indicated as in **a. c** The distributions of overlap fractions for mask pairs grouped using the fuzzy size threshold described in **a**. We omit pairs that do not overlap for clarity. **d** Applying morphological erosions of the cell masks reduces the number of overlapping cell pairs, but generates smaller masks. We judge masks with areas below 10 pixels squared to be too small. **e** The numbers of overlapping cell pairs remaining from the training, validation and test sets either before (denoted None) or after splitting into size categories and applying an optimised number of erosions.

We therefore choose the size categories so that most overlaps occur between pairs of cells in different categories and little overlap occurs between pairs of cells within a category. For example, rather than converting the cell masks directly into a single binary image for training, if first we divide cells into two size categories and convert the masks within each category to a separate binary image, giving two images rather than one, then in these two images we have eliminated all overlaps occurring between cells in the smaller category with cells in the larger category (Appendix 3 Fig. 1 & 2a – lower triangle).

To divide the cell masks into two categories, we define a fuzzy size threshold using a threshold *T* and padding value *P*. The set of smaller masks is all masks whose area is less than *T* + *P* ; the set of larger masks is all masks whose area is greater than *T* − *P*. Consequently, it is possible to have the same mask in both sets (Appendix 3 Fig. 1c). This redundancy ensures the CNN can produce confident predictions even for cells close to the size threshold – we eliminate any resulting duplicate predictions in post-processing. A pair of masks are prevented from overlapping, by being converted into distinct binary images, if their sizes are separated by the threshold (after padding): the two masks must be in different size categories, and so the smaller cell must have a size *<T* − *P* and the larger cell must have a size *>T* + *P*. To scale with pixel size, we set *P* to be 3% of the area of the largest mask in the training set.

To determine an optimal fuzzy threshold, we test 29 values evenly spaced between the minimal and maximal mask sizes and choose the threshold that minimises the summed overlap fraction for all mask pairs not excluded by the threshold. Even with just one fuzzy threshold (Appendix 3 Fig. 2a), most of the pairs with substantial overlap – typically buds with neighbouring cells – are excluded (Appendix 3 Fig. 2c).

After applying the threshold, overlaps between cells within a size category remain, and we reduce such overlaps using morphological erosions (Appendix 3 Fig. 1). We use the training data to optimise the number of erosions applied to each size category. As the number of iterations increases, there is a trade-off between the number of overlapping mask pairs and the number of masks whose eroded areas become too small to be confidently predicted by the CNN (Appendix 3 Fig. 2d). Without erosion, many of the large cells show overlaps; with too much erosion, the smallest masks distort their shapes or disappear completely. We therefore optimise the number of iterations separately for each size category, picking the highest number of iterations that can be applied without any of the training masks in that size category falling below an absolute minimal size, defined as 10 pixels squared.

Combining categorising by size with eroding reduces the number of pairs of overlapping masks almost to zero (Appendix 3 Fig. 2e). We arrive at three size categories by first introducing an additional fuzzy threshold for each of the two initial size categories. These thresholds are similarly determined by testing 29 fuzzy threshold values and calculating the overlap fraction for all mask pairs not excluded by either the original or the new threshold. We only keep one of the new thresholds – the one minimising the overlap fraction, giving three size categories in total. This extra category results in a further, although proportionally smaller, decrease in the number of overlapping masks.

After erosion, mask interiors within each size category are easily identified, but with less resolved edges. To help alleviate this loss, we generate edge targets from the training data (Appendix 3 Fig. 1e) – the outlines of all cells within each size category.

#### Three types of training targets

The curated data is further annotated with lineage assignments (Appendix 3 Fig. 1f), which we use to generate ‘bud neck’ targets (Appendix 3 Fig. 1g).

In total, then, the training targets for the CNN are the mask interiors and the edges for three size categories and the bud necks.

#### Predicting semantic targets with a convolutional neural network

We trained fully convolutional neural networks [68] to map a stack of bright-field sections to multiple binary target images. Example inputs and outputs are shown in Appendix 3 Fig. 1, but we also trained networks with only one or three bright-field sections. The intensities of the bright-field sections were normalised to the interval [− 1, 1] by subtracting the median and by scaling according to the range of intensities expected between the 2^nd^ and 98^th^ percentiles.

Each output layer of the CNN approximates the probability that a given pixel belongs to the target class, being a convolution with kernel of size 1 × 1 and sigmoidal activation. All other convolutions had kernels of size 3 × 3 with ReLU activation and used padding to ensure consistent dimensions for input and output layers. Different core network architectures were tested and trained.

#### Augmenting the training data

To prevent over-fitting and improve generalisation, we augmented the training data [68]. Each time a training example was presented to the CNN, a randomly selected series of image manipulations was applied to the input and target. The same training example therefore typically appears differently for each epoch.

One augmentation was always applied and the others applied with a certain probability. If the bright-field input had more Z sections than expected by the network, we selected a random subset, excluding any subsets with selected sections that were separated by two or more missing sections. The remaining augmentations were applied with a probability *p* and comprised vertical and horizontal translation (each with *p* = 0.25), image rotation (*p* = 0.2), rescaling (*p* = 0.08), vertical and horizontal flips (each with *p* = 0.08), addition of white noise (*p* = 0.08), and a step shift of the Z sections (*p* = 0.08). The probability of not augmenting was thus *p* = 0.30. To show a different region of each image-mask pair at each epoch, translation, rotation and re-scaling were all applied to images and masks before cropping to a consistent size (96 96 pixels for a pixel size of 0.182 *μ*m). Using reflection to handle the boundary, translations were over a random distance and rotations over a random angle. The factor used for re-scaling was randomly adjusted by up to 5%. Augmentation by addition of white noise involved adding random Gaussian noise with a standard deviation picked from an exponential distribution with rate 0.003 to each pixel of the (normalised) bright-field images.

To reduce aliasing errors when manipulating binary masks during augmentation, all image transformations were independently applied to each filled mask before converting the transformed masks into one binary image. Further, before a transformation, each binary filled outline was smoothed with a 2D Gaussian filter, and after the transformation, the transformed binary outline was found by the Canny algorithm. To determine the standard deviation of this Gaussian filter, σ, we tested a range of values on the training outlines. For each filled outline and σ,we applied the filter followed by edge detection and filling. We then calculated the intersection over union of the resulting filled outline with the original filled outline. We observed that as a function of edge length, defined as the number of pixels in the outline, the *σ* producing the highest intersection over union increased exponentially. We consequently used an exponential fit of this data to estimate an appropriate *σ* for each outline.

**Figure 3:**
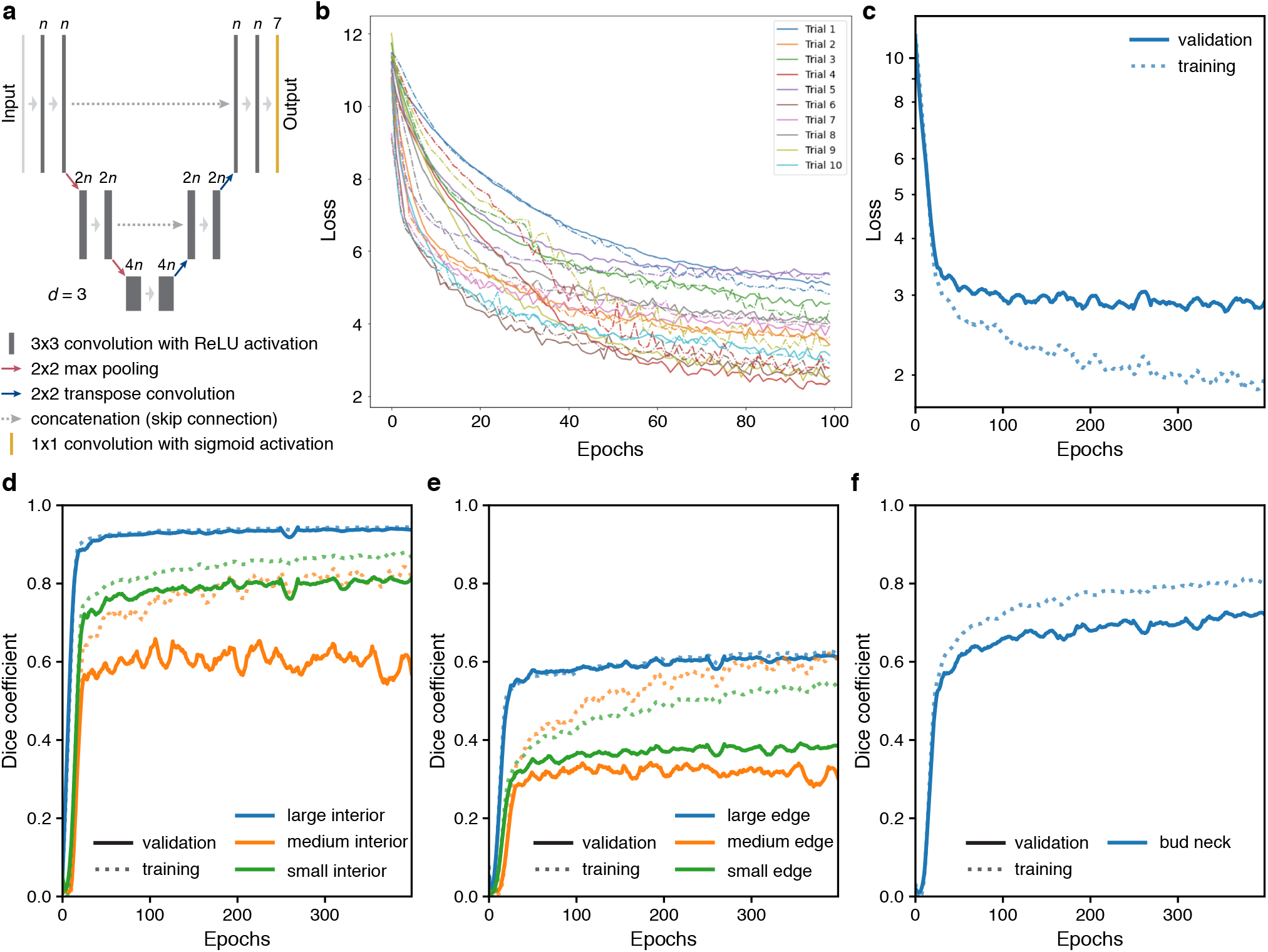
Training performance of the multi-target U-Net. **a** Schematic U-Net architecture with depth *d* = 3. Labels above convolution operations indicate number of output filters as a multiple of *n*. Layer heights indicate reduction in image width/height with network depth. **b** Training (solid) and validation (dot-dash) loss for 10 training trials using the U-Net architecture each with randomly chosen hyperparameters. **c** Loss for the fully trained U-Net with hyperparameters chosen from the trial network with the lowest final validation loss (a U-Net with depth *d* = 4, filter factor *n* = 8 and batch normalisation). **d–f** Performance of (d) interior, (e) edge and (f) bud neck targets by the U-Net of **c** decomposed into the three different size categories when possible. The Dice coefficient reports similarity between prediction probabilities and target masks with a value of 1 indicating identity. For two sets *X* and *Y*, the Dice coefficient is <u>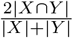</u>.

#### Training

Networks were trained using Keras with TensorFlow 1.14. We used Adam optimisation with default parameters except for a learning rate of 0.001 and regularised by keeping only the network weights from the epoch with the lowest validation loss (similar in principle to the early stopping method) [68]. We train for 400 epochs, or complete iterations over the training data set.

The loss function is the sum of the binary cross-entropy and one minus the Dice coefficient across all targets:

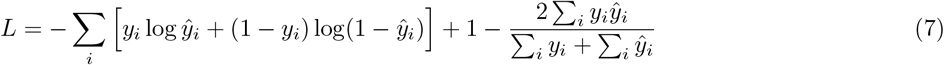

where *y* is the tensor of true values, ŷ is the CNN’s sigmoid tensor output of the CNN, and *i* is a vectorised index.

Each CNN is trained to a specific pixel size, and we ensured that training images and masks with different pixel sizes were re-scaled appropriately

#### CNN architectures

We trialled two core architectures for the CNN – U-Net [72] (Appendix 3 Fig. 3a) and Mixed-Scale-Dense (MSD) [73] – and optimised hyperparameters to find the smallest loss on the validation data.

The U-Net performed best (see ‘Optimising hyperparameters’ below). The U-Net has two parts: an encoder that reduces the input into multiple small features and a decoder that upscales these features into an output [72]. Each step of the encoder comprises a convolutional block, which creates a new, larger set of features from its input. To force the network to keep only small, relevant features, a down-sampling step is applied after three convolutional blocks. This maximum pooling layer reduces the size of the features by half by replacing each two-by-two block of pixels by their maximal value. The decoder also comprises convolutional blocks, but with up-sampling instead of down-sampling. The up-sampling step is the inverse of down-sampling: each pixel is turned into a two-by-two block by repeating its value. Finally, most characteristic of the U-Net is its skip layers. These layers preserve information on the global organisation of the pixels by passing larger-scale information from the encoder to the decoder after each up-sampling step. They act by concatenating the same-size layer of the encoder into the decoder layers, which are then used as inputs for the next step of the decoder. The decoder is therefore able to create an output from both the local features that it upsampled and from the global features that it obtains from the skip layers.

For the U-Net, we optimised for depth, for a scaling factor for the number of filters output by each convolution, whether or not to include batch normalisation, and for the proportion of neurons to drop out on each batch. For the MSD, we optimised for depth, defined as the total number of convolutions, for the number of dilation rates to loop over, with each loop increasing dilation by a factor of two, for an overall dilation-rate scaling factor, and whether or not to include batch normalisation.

#### Optimising hyperparameters

We used `KerasTuner` with TensorFlow 2.4 to optimise hyperparameters (Appendix 3 Fig. 3b), choosing random search with default settings, training for a maximum of 100 epochs, and having 10 training and validation steps per epoch. The U-Net and MSD networks with the lowest final validation loss were then re-trained as described (Appendix 3 Fig. 3c), and the network with the lowest validation loss chosen.

For our data, the best performing model was a U-Net with depth four, and so three contractions, with a scaling factor of eight for the number of filters output by each convolution, giving 8, 16, 32 and 64 filters for each of the two chained convolution layers of the encoding and decoding blocks, with batch normalization, and with no drop-out. Its performance is shown in Appendix 3 Fig. 3d–f.

#### Identifying cells

To identify cell instances from the semantic predictions of the CNN, we developed a post-processing pipeline (Appendix 3 Fig. 4a). Post-processing can be split into two parts: proposing unique cell outlines and refining edges.

The pipeline includes multiple parameters that we optimise on validation data by a partial grid search. We favour precision (the fraction of predicted positives that are true) over recall (the fraction of ground truth positives that are predicted) by maximising the *F*_*/3*_ score with, *B* = 0.5. Recall that for true positives TP, false negatives FN, and false positives FP,

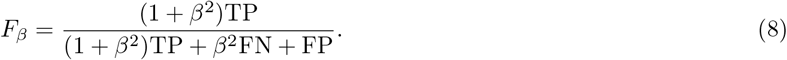

We measure how well two masks match using the intersection over union (IoU) and consider a match to occur when the IoU *>* 0.5. Nevertheless, multiple predictions may match a single target mask because predicted masks can overlap too. We therefore count true positives as target masks for which there is at least one predicted mask with IoU *>* 0.5. Any predicted masks in excess of the true positive count become false positives, thus avoiding double counting. Unmatched target masks are false negatives.

#### Proposing cell outlines

The post-processing pipeline starts by identifying candidate outlines independently for each size category. The CNN’s outputs are images 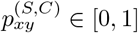 approximating the probability that a pixel at position (*x, y*) belongs to either the small, medium, or large size categories, denoted *S*, and to either the interior (Appendix 3 Fig. 4b) or edge (Appendix 3 Fig. 4e & f) target classes, denoted *C*.

In principle, we could find instances for each size category by thresholding the interior probability 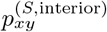 and identifying connected regions as outlines. To further enhance separability, however, we also reweight the interior probabilities using the edge probabilities. Specifically, we identify connected regions from semantic bitmasks 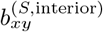 by those pixels that satisfy

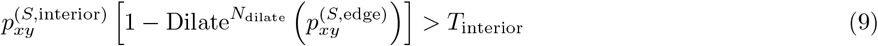

where Dilate^*N*^ specifies *N* iterations of a grayscale morphological dilation and *T*_interior_ is a threshold. We optimise the thresholds *T*_interior_ [0.3, 0.95], number of dilations *N*_dilate_ ϵ {0, 1, 2 }, and the order of connectivity (1- or 2- connectivity) for each size category.

**Figure 4:**
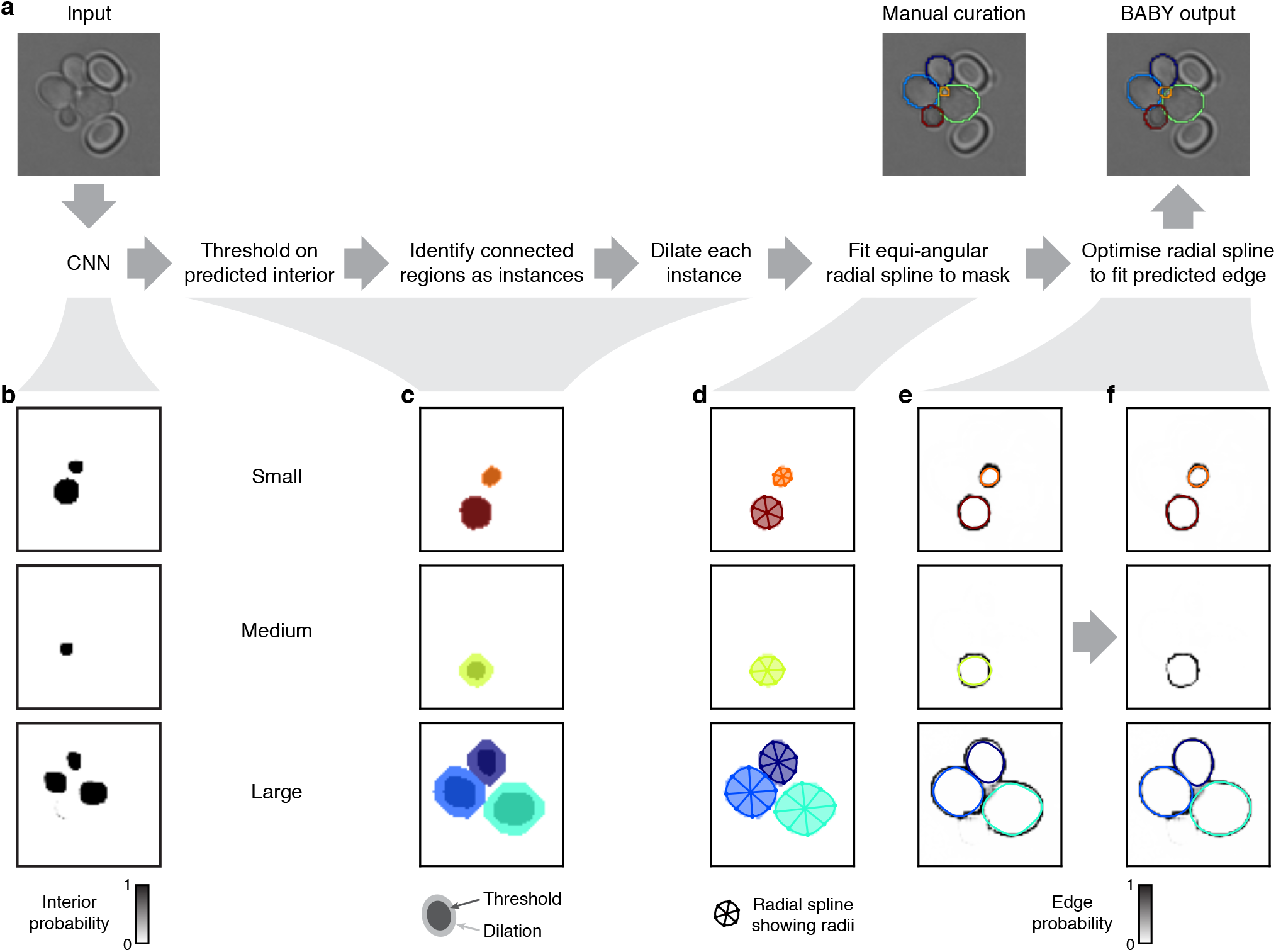
Segmenting overlapping cell instances from the CNN output. **a** Flow chart summarising the post-processing for identifying individual instances from the CNN’s multi-target output. Here and below, we show results using the five Z sections of Appendix 3 Fig. 1 as input for the CNN. One of which is repeated here. **b** Probability maps output by the CNN for the interiors of small, medium and large cells. **c** Bitmasks obtained by thresholding on the CNN’s output. Darker shading shows bitmasks before each instance is dilated to compensate for the erosion applied when generating the training targets. Colour indicates distinctly identified instances. **d** The initial, equiangular radial splines proposed for each instance overlaid on the dilated bitmasks from **c**. The rays defining placement of the knots are shown as spokes. **e** The same initial proposed radial splines overlaid on the edge target probability maps output by the CNN. **f** The radial splines after optimisation to match edge probabilities. The outline in the medium size category is detected as a duplicate and is not optimised.

The connected regions in 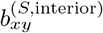 define masks that are initial estimates of the cells’ interiors (darker shading in Appendix 3 Fig. 4c). The actual cell interiors used to train the CNN are generated by iterative binary morphological erosions of the full mask, where the number of iterations *N*_erosion_ is pre-determined for each size category. First, we remove small holes and small foreground features by applying up to two binary morphological closings followed by up to two binary morphological openings. Second, we estimate full masks 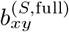 from each putative mask by applying *N*_erosion_ binary dilations (light shading in 4c), undoing the level of erosion on which the CNN was trained. Both the numbers of closing, *N*_closing_, and opening, *N*_opening_, operations are optimised.

Masks whose area falls outside the limits for a size category are discarded. For each category, however, we soften these limits, on top of the fuzzy thresholds, by optimising an expansion factor *F*_exp_ ϵ [0, 0.4], which extends the limits by a fractional amount of that category’s size range. We also optimise a single hard lower threshold *T*_min_ ϵ [0, 20] on mask area.

#### Using splines to describe mask edges

To prepare for refining edges and to further smooth and constrain outlines, we use a radial spline to match the edge of each of the remaining masks (Appendix 3 Fig. 4d). As in DISCO [30], we define radial splines as periodic cubic B splines using polar coordinates whose origin is at the mask’s centroid.

We generalise this representation to have a variable number *n*^(*S,i*)^ of knots per mask specified by *n*^(*S,i*)^-dimensional vectors of radii ***r***^(*S,i*)^ and angles θ^(*S,i*)^:

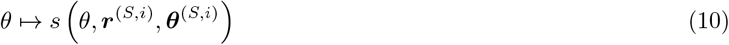

A mask’s outline is then fully specified by those pixels that intersect with this spline.

To initially place the knots, we search along rays originating at the centroid of each mask 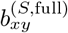 and find where these rays intersect with the mask edge. We determine the edge by applying a minimum filter with 2-connectivity to the mask and set to true all pixels in the filtered image that are different from the original one. We then smooth the resulting edge image using a Gaussian filter with *σ* = 0.5. For a given polar angle θ, we find the radius of the corresponding knot by averaging the edge pixels that intersect with the ray, weighted by their values. We use the major axis of the ellipse with same normalized second central moment (`regionprops` function from Scikit-image [71]) as the mask to determine both the number of rays, and so knots, and their orientations. The length ℓ^(*S,i*)^ of the major axis gives the number of rays: four for 0 *<* ℓ^(*S,i*)^ *<* 5; six for 5 ℓ^(*S,i*)^ *<* 20; and eight for ℓ^(*S,i*)^ 20. For this initial placement, we choose equiangular θ^(*S,i*)^, with the first knot on the ellipse’s major axis.

#### Discarding poor or duplicated outlines

The quality of the outline masks 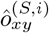 derived from these initial radial splines are then assessed against the edge probabilities generated by the CNN (Appendix 3 Fig. 4e) and masks of poor quality discarded. We calculate the edge score for a given outline as

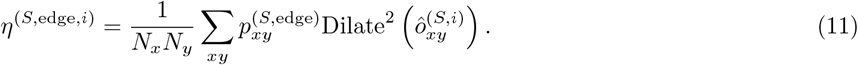

We discard those outlines for which the edge score is less than a threshold, where the thresholds *T*_edge_ ϵ [0, 1) are optimised for each size category based on the range of edge scores observed.

With a smoothed and filtered set of outlines, we proceed by detecting and eliminating any outlines duplicated between size categories. We start by filling the outlines to form a set of full masks 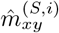. We then compare these masks between neighbouring size categories *S*_*j*_ and *S*_*k*_. We consider the pair of masks *i*_1_ and *i*_2_ to be duplicated if one of the masks is almost wholly contained within the other:

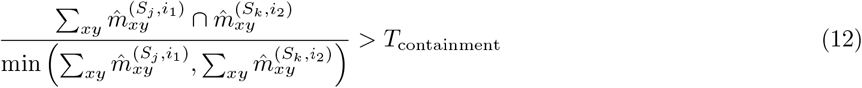

for some threshold *T*_containment_ ϵ [0, 1], optimised on validation data. For pairs that exceed this threshold, we keep only the mask with the highest edge score, Eq. 11.

For each size category, the first part of the post-processing pipeline finishes with the set of outlines that pass these size, edge probability, and containment thresholds. Appendix 3 Table 1 gives values for the optimised post-processing parameters.

**Table 1:** Optimised post-processing parameters for standard model. The standard model takes five brightfield Z sections with a pixel size of 0.182 *μ*m as input. Excepting *T*_min_ and *T*_containment_, parameters are optimised separately for each size category.

| Parameter | $S = \text{small}$ | $S = \text{medium}$ | $S = \text{large}$ |
| --- | --- | --- | --- |
| $T_{\text{interior}}$ | 0.35 | 0.5 | 0.95 |
| $N_{\text{dilate}}$ | 0 | 0 | 0 |
| Connectivity | 2 | 1 | 1 |
| $N_{\text{closing}}$ | 0 | 0 | 0 |
| $N_{\text{opening}}$ | 0 | 0 | 2 |
| $F_{\text{exp}}$ | 0.32 | 0.06 | 0.28 |
| $T_{\text{edge}}$ | 0.0012 | 0.0028 | 0.0 |
| $T_{\text{min}}$ | | 19 | |
| $T_{\text{containment}}$ | | 0.85 | |

##### Refining edges

The outlines 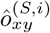, defined by the radial splines, do not directly make use of the CNN’s edge targets for their shape and deviate from 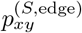, particularly for those in the large size category (Appendix 3 Fig. 4e).

We therefore optimise the radial splines to better match the predicted edge. This optimisation is challenging because 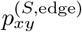 provide only a semantic representation of the edge – the association of a given pixel (*x, y*) with a particular instance *i* is unknown. Our approach is to use the outlines to generate priors on whether predicted edge pixels associate with a given instance. We then apply standard techniques to optimise the fit of the radii and angles of the knots for each outline’s spline to its instance’s likely edge pixels.

**Figure 5:**
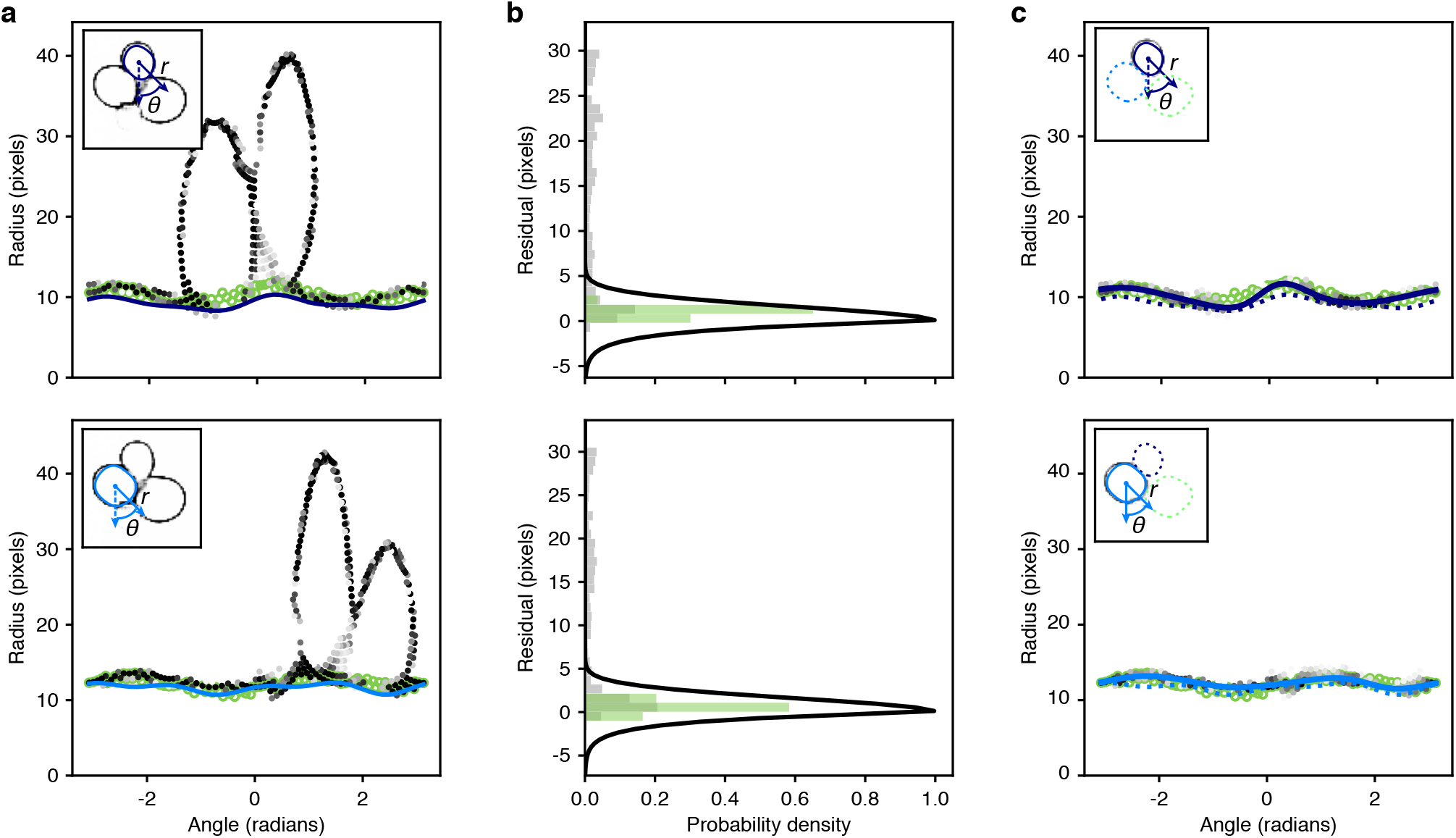
Optimisation of the radial spline to fit the predicted edge. **a** Predicted edge pixels by the CNN in polar coordinates centred on two different instances (top and bottom panels). Darker shading indicates a higher probability of being an edge. Open green circles are the manually curated ground truth. Solid lines are the initial radial splines estimated from the CNN’s predictions of interiors. Insets show the predicted edge in cartesian coordinates with the instance providing the origin marked by its initial outline and the indicated polar coordinates. **b** Binned residuals of the predicted edge pixels with the initial radial spline for the examples shown in **a**. Only edge pixels with probability greater than 0.2 are considered. Binned residuals for the ground truth are in green. Black lines show the function used to re-weight pixel probabilities for each instance. **c** As for **a**, but after the edge pixels have been re-weighted for each instance. Solid lines indicate the optimised radial spline. The inset shows with a solid line the outline favoured by the instance-association probability and the dashed lines those outlines disfavoured.

To associate pixels with instances, we first calculate the radial distance of each pixel from the initial radial spline function *s* proposed for an instance, 10. To increase speed, we consider only pixels where 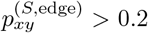. Expressing the edge pixels in polar coordinates as (ρ, ϕ) with the origin at the instance’s centroid, this distance is

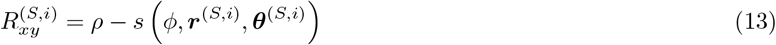

which we will refer to as a pixel’s residual. We give two examples of edge pixels (Appendix 3 Fig. 5a) and of the corresponding residuals (Appendix 3 Fig. 5b), which highlight the need to associate pixels with a given instance before attempting to optimise the spline.

We use the residuals, Eq. 13, to assign prior weights to pixels:

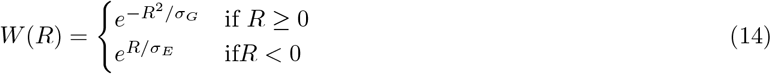

] where *σ*_*G*_ = 5 and *σ*_*E*_ = 1. The function *W* is a Gaussian function of the residual for pixels exterior to the proposed outline and an exponential function for pixels interior (Appendix 3 Fig. 5b). This asymmetry is intended to increase tolerance for interior edge pixels, which may belong to neighbouring instances that overlap with the cell of interest. In such cases, instance association should be improved, particularly where the edges of each of the cells intersect.

With these prior weights, we find the probability that each edge pixel is associated with a particular instance and not with the others, via:

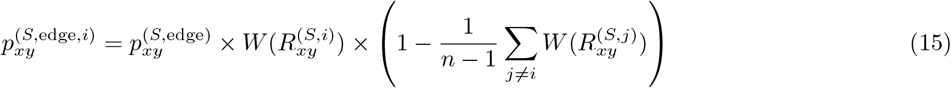

where *n* is the number of detected instances in this and adjacent size categories, with *j* running over all these instances. We filter the result, keeping only those edge pixels with *p*^(*S*,edge,*i*)^ *>* 0.1. Examples are shown in Appendix 3 Fig. 5c. We optimise the knot radii ***r***^(*S,i*)^ and angles θ^(*S,i*)^ for each radial spline by minimising the squared radial residual between the spline and the edge pixels, Eq. 13. Residuals are weighted by 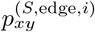, Eq. 15, and initial values are taken from each 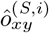. Radii are constrained to a 30% change from their initial values; angles are constrained to a change of 25% of the initial angular separation between knots: *θ*_*i*+1_ *θ*_*i*_. The resulting optimised radial spline provides the outlines output by the BABY algorithm.

## Appendix 4

### The BABY algorithm: tracking cells and identifying lineages

To track cells and lineages, we have two tasks: first, link cell outlines from one time point to the next (Appendix 4 Fig. 4a), and second, identify mother-daughter relationships (Appendix 4 Fig. 4b).

**Figure 1:**
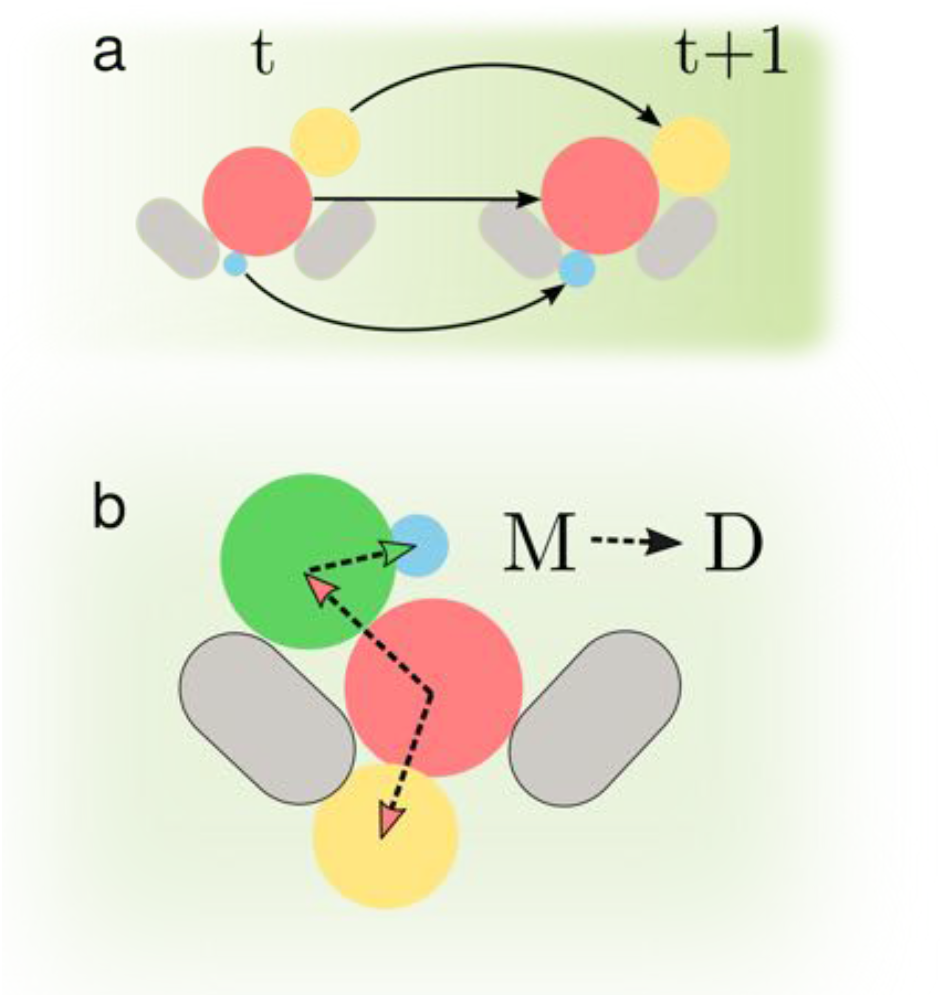
Determining accurate lineages requires solving two independent tasks. **a**. We must identify cells across time points regardless of how they grow and move within the images. **b**. We have to find the mother-daughter relationship between cells at every time point.

### Tracking cells from image to image

In our setup, daughter cells may be washed out of the microfluidic device and so disappear from one time point to the next. These absences undermine other approaches to tracking, such as the Hungarian algorithm [29].

To track cells, we use the changes in their masks over time to indicate identity. From each mask, we extract an array of attributes, such as the mask’s area, major axis length, etc., and to compare a mask at one time point to a mask at another time point, we subtract the two corresponding arrays of features. This array of differences is the array of features for the classification algorithm.

Our training data comprises a series of manually labelled time-lapse images from four experiments. For two consecutive time points, we calculated the difference in feature arrays between all pairs of cells and grouped these difference arrays into two classes: one for identical cells – cells with the same label – and one for all other cells.

### Using multiple time points in the past

To generate additional training data, we use multiple time points backwards in time. For example, for time *t*, we generate not only feature vectors by comparing with cells at *t* 1, but also with cells at *t* − 2 and *t* − 3. We found this additional data increased generalisability, allowing accuracy to be maintained across a wider range of imaging intervals and growth rates. For the purpose of training, we treat the additional data as consecutive time points – the algorithm does not know whether the features come from one or more than one time point in the past.

As part of testing that all features contribute to the learning, we divided the features into two overlapping sets. One set had no features that explicitly depend on distance, comprising area, lengths of the minor and major axes, convex area, and area of the bounding box; the second set did include distance-dependent features, comprising area, lengths of the minor and major axes, and convex area again, but additionally including the mask’s centroid, and the distance obtained from the *x*- and *y*-axis locations.

We compared three standard algorithms for classification [74]: the Support Vector Classifier (SVC), Random Forest, and Gradient Boosting, specifically Xtreme Gradient Boosting [75]. We used `scikit-learn` [70] to optimise over a grid of learning hyperparameters.

For the SVC, we considered a regularisation parameter *C* of 0,1, 10, or 100; a r kernel coefficient of 1, 10 ^− 2^, or 10 ^− 4^; no shrinking heuristic to speed up training; and either a radial basis function or sigmoid kernel.

For the Random Forest, we explored a range between 10 and 80 estimators and a depth between 2 and 10 levels.

For Gradient Boosting, we used a maximal depth of either 2, 4, or 8 levels; a minimal child weight of 1, 5, or 10; gamma, the minimal reduction in loss to partition a leaf node, of 0.5, 1, 1.5, 2, or 5; and a sub-sampling ratio of 0.6, 0.8, or 1.

**Figure 2:**
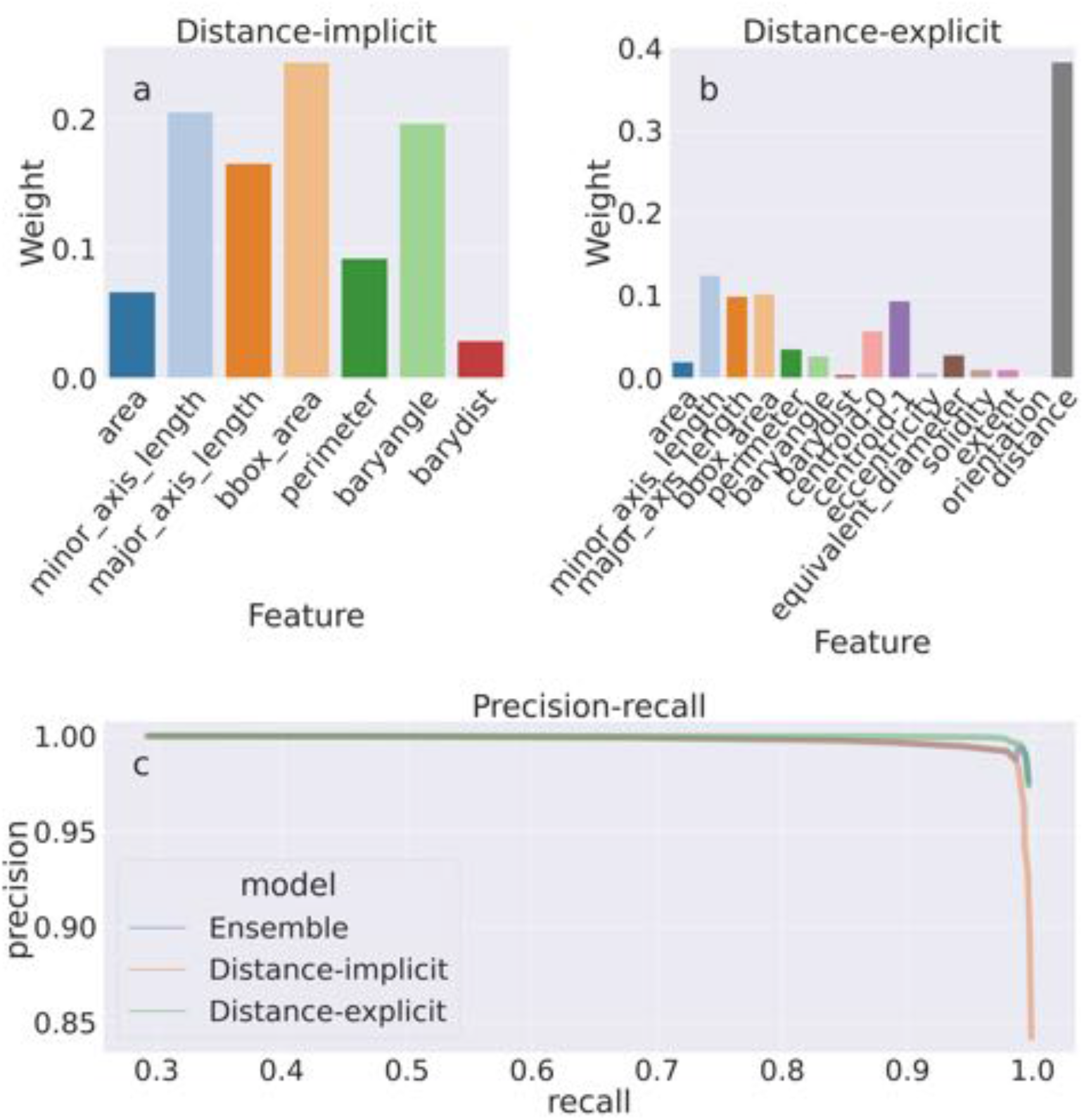
The importance of the features used by the Random Forest classifier for tracking cells from time point to time point. Depending on the features with which we train the classifier, the weights, or feature importances, are more evenly spread. **a** If we train the classifier using features that explicitly include distance-dependence, distance drives the decisions, and the remaining features are only used for marginal cases. **b** If we train the classifier using distance-implicit features, however, the weights are more uniform. **c** The precision-recall curve shows high accuracy for both sets of features.

Within the training data, the number of time points for each experiment is different and the feature arrays will favour linking cells within an experiment. To prevent biases toward long experiments, we define the accuracy as the fraction of true positives – cells correctly linked between images, and compare the precision and recall of this time-averaged accuracy.

After training, we evaluated which features are important for classifying, using the Random Forest. The distribution of the features’ importance depends on whether distance is included (Appendix 4 Fig. 2): excluding distance distributes the weight of decision more evenly.

### An ensemble model

The precision-recall curve indicates that using the distance-explicit features is best, although both sets of features have high accuracy (Appendix 4 Fig. 2c). Although performing better on our test data, we expect that using the distance-explicit features may perform worse if the cells pivot or are displaced in some other way. Therefore, we use the non-explicit features as our main model, but also use the distance-explicit features to resolve any ambiguous predictions. The ensemble model performs similarly to the distance-implicit classifier, but for more stringent thresholds becomes behaves like the distance-explicit model.

### Making predictions

To predict with the classifier, we use data from the current time point and the two most recent previous time points. We generate feature arrays between *t* and independently *t* − 1 and *t* − 2 and feed both arrays to the classifier. If the probability returned is greater than 0.9, we accept the result; if the probability lies between 0.1 and 0.9, we use instead the probability returned by the backup classifier, which uses the distance-explicit features.

Using multiple time points to track cells has two advantages: first, it reduces noise when there is an artefact, either in image acquisition, such as a loss of focus, or in segmentation; second, it ensures that cells are more consistently identified if their position or shape transiently changes. Including data further back in time is neither computationally efficient nor more accurate – greater than three time points is over 15 minutes in our experiments, about a sixth of a cell cycle.

We apply the linear sum assignment algorithm, via `SciPy`, on the probability matrix of predictions to assign labels (Appendix 4 Algorithm 1), which guarantees at most one outline assigned to each cell by choosing the set of probabilities whose total sum is highest. To match a cell with its previous self, we pick the cell in the recent past that generates the highest probability when paired with the cell of interest, providing this probability is greater than 0.5. A cell is labelled as new if the probabilities returned from pairing with all cells in the recent past is below 0.5.

#### Algorithm 1: Cell labelling

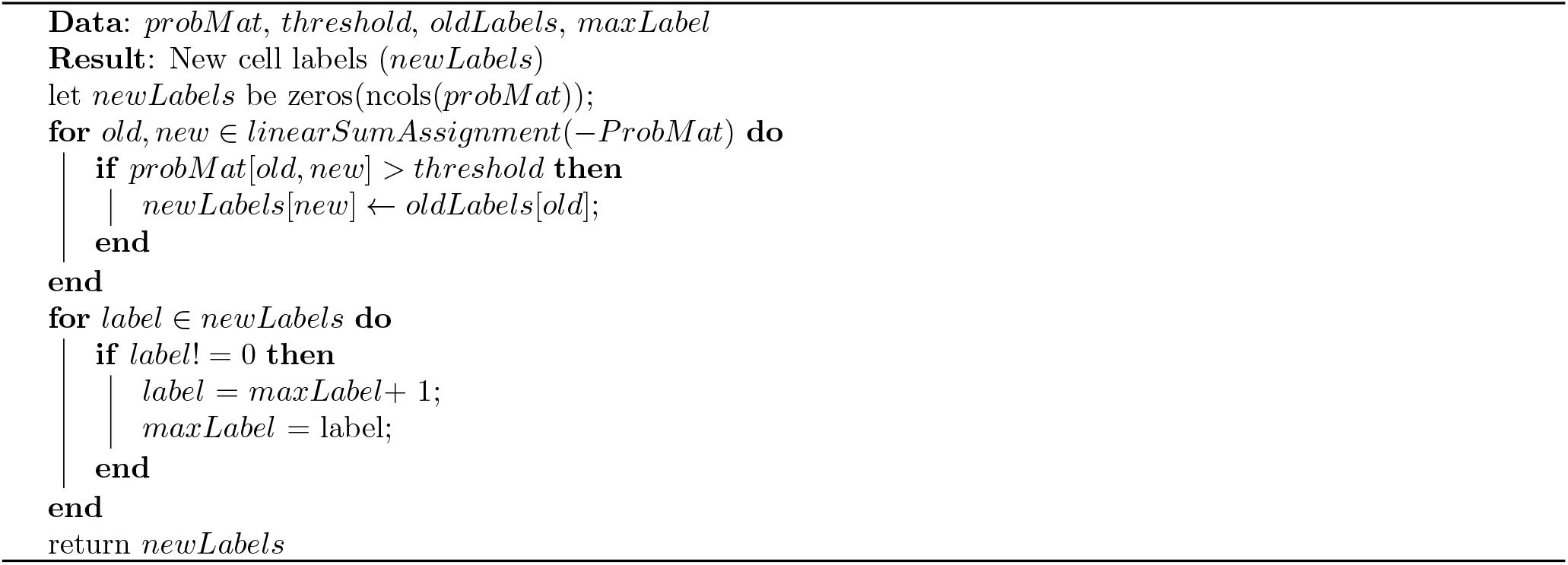

### Assigning lineages

We wish to identify which cells are buds of mothers and which mothers have buds. This problem is analogous to tracking, but, rather than identifying pairs of cells that are the same cell at different time points, we must identify pairs of cells that are a mother-bud pair at one time point. We therefore seek to determine the probability that a pair of cells is a mother-bud pair (Appendix 4 Fig. 3). Unlike for tracking, however, we anticipated that the cell outlines alone would be at best a weak indicator of this probability.

### Defining mother-bud features

We observed that cytokinesis is sometimes visible in bright-field images as a darkening of the bud neck and designed features to exploit this characteristic of mother-bud pairs.

Multiple of these features rely on the CNN’s prediction of bud necks. For generalisability and to avoid ambiguity, we chose to define the corresponding training target using manually annotated outlines and lineage relationships, rather than relying on a fluorescent bud-neck marker. Specifically, we define a binary semantic ‘bud-neck’ training target that is true only at pixels where a mother mask, dilated twice by morphological dilation, intersects with its assigned bud (Appendix 3 Fig. 1). Assigning a time of cytokinesis by eye is challenging, and so we included two constraints to identify a bud. First, the bud must be current – as soon as another bud is found associated to the mother, the previous bud is excluded. Second, buds are excluded if their area is larger than 10 *μ*m^2^ (300 pixels for our standard training target with a pixel size of 0.182 *μ*m and corresponding to a sphere of ∼ 24 *μ*m^3^).

**Figure 3:**
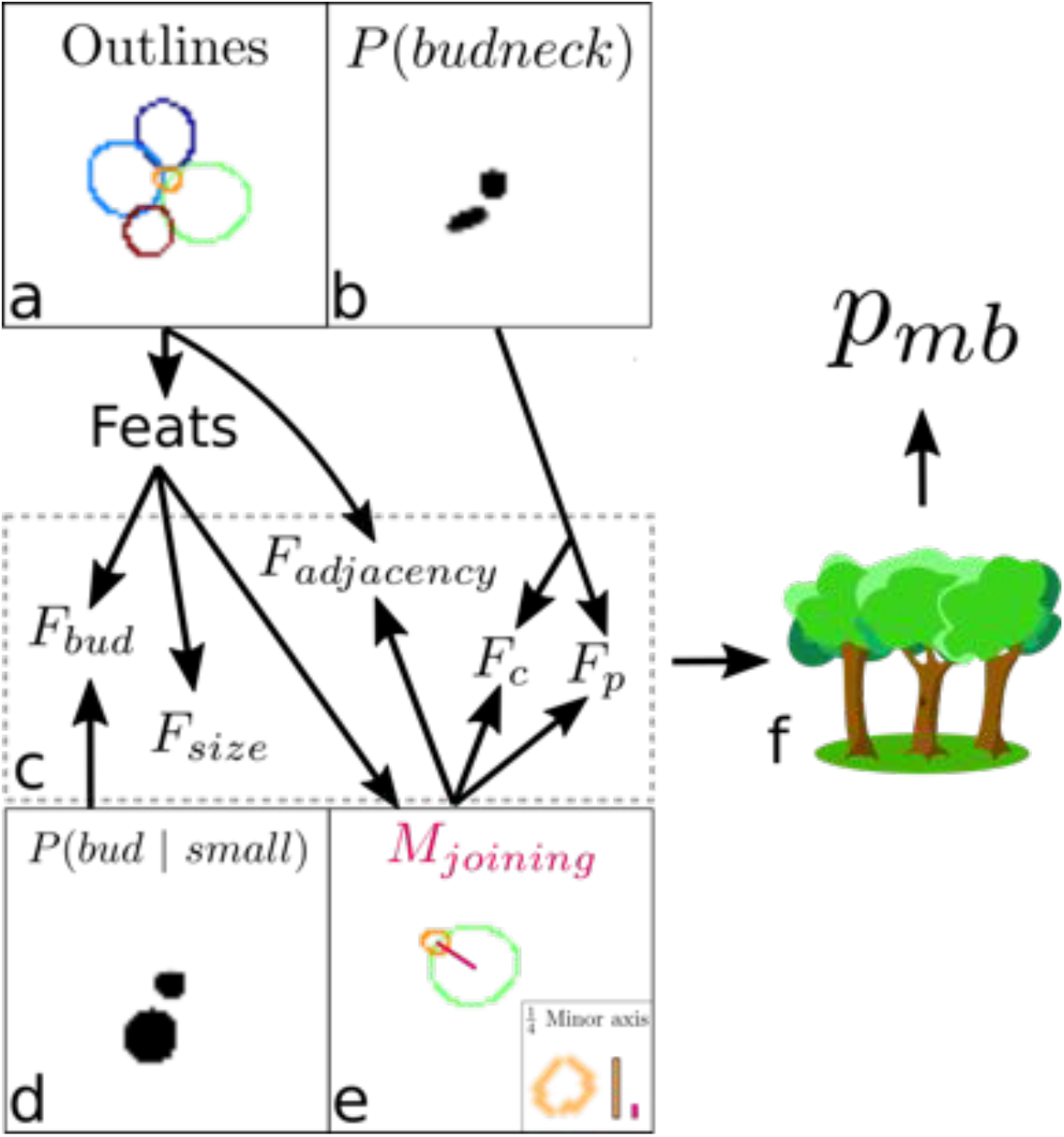
Overview of the algorithm for assigning lineages. **a** Cell outlines. Colours are only illustrative. **b** The CNN’s predicted probabilities per pixel of being a bud neck are shown for small cells. White is zero, and different colour intensities are probabilities. **c** Composite features used by the classifier to solve the task. **d** The probability of small cells being a bud. **e** An intermediate element of assigning lineages is defining *M*_joining_ – the red rectangular box. **f** Feeding the features into a trained random forest model returns the probability of a pair of cells being a mother and bud.

We used five image features to characterise a mother-bud relationship. For an ordered pairing of all cells in an image, we consider a putative mother-bud pair and define a mask, *M*_joining_, as the joining rectangle between the centres of the mother and bud with a width equal to one quarter the length of the bud’s minor axis. Given *M*_joining_, the features are:

i. *F*_size_, which is the ratio of the mother’s to bud’s area. Mothers generally have a greater size than their bud so that *F*_size_ *>* 1.
ii. *F*_adjacency_, which is the fraction of *M*_joining_ intersecting with the union of the mother’s and bud’s masks. Mothers should be proximal to their buds so that *F*_adjacency_ is close to one – only a small fraction of *M*_joining_ should lie outside of the mother and bud outlines.
iii. *F*_bud_, which is the mean over the union of the CNN’s output for a small, interior targets and all pixels contained in the bud. *F*_bud_ approximates the probability that a cell is a bud and should be close to one for mother-bud pairs.
iv. *F*_p_, which is the mean over the union of the CNN’s output for bud-necks, only including those pixels whose probability is greater than 0.2, with the pixels contained in *M*_joining_. *F*_p_ approximates the probability that the mother and bud are joined by a bud neck.
v. *F*_c_, which is the number of the CNN’s bud-neck target pixels that have a probability greater than 0.2 and are in *M*_joining_ normalised by the square root of the bud’s area, or effectively the bud’s perimeter. We interpret *F*_c_ as a confidence score on *F*_p_ because a single spurious pixel with high bud-neck probability could produce high *F*_p_.

### Training a classifier

With these features, we train a random forest classifier to predict the probability that a pair of cells is a mother and bud. We train on all pairs of cells in the validation data to ensure that the classifier does not rely on the CNN’s high performance on the training data. We optimised the hyperparameters, including the number of estimators and tree depth, using a grid search with five-fold cross-validation. We optimise for precision because true mother-bud pairs are in the minority and because our strategy for assigning lineages aggregates over multiple time points (as detailed below).

For our standard CNN trained to a pixel size of 0.182 *μ*m^2^, the random forest classifier had a precision of 0.80 and recall of 0.47 on the test data. This data has 211 true mother-bud pairs out of 1654 total pairs. The classifier assigned feature importances of 0.42 for *F*_size_, 0.10 for *F*_adjacency_, 0.26 for *F*_bud_, 0.08 for *F*_p_, and 0.13 for *F*_c_.

### Assigning each cell a unique mother

To establish lineage relationships, we need to assign at most one tracked mother cell to each tracked cell object. We use the classifier to assign a mother-bud probability 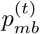 for each time point at which a pair of tracked objects are found together. We then estimate the probability that a tracked object ĉ has ever been a mother using

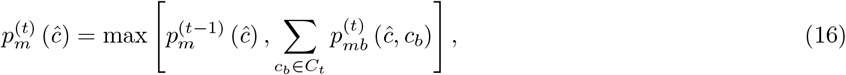

as well as the probability that it has ever been a bud with

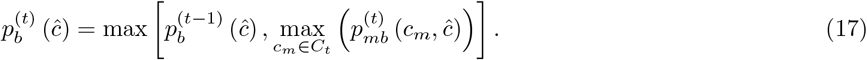

Finally, for a putative mother-bud pair we calculate a cumulative score that is reduced if the candidate bud has previously shown a high probability of being a mother:

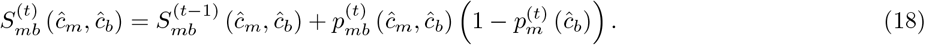

At each time point, we then propose lineages by assigning each putative cell object ĉ_*b*_ with a bud probability 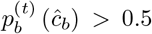 and a mother 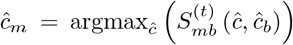. We treat the mother-bud assignments proposed at the final time point as definitive, because they have integrated information over the entire time series, and avoid spurious assignments by requiring all buds to be present for at least three time points.

#### Post-processing

Though rare, we do have to mitigate occasional detection, tracking and assignment errors. For example, debris can occasionally be mistakenly identified as a cell and tracked.

We discard tracks that have both small volumes and show limited growth over the experiment. Specifically, we discard a given cell track *i* with duration *T*_*i*_, minimal volume 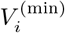 at time 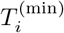, and maximal volume 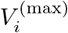 at time 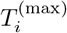 if both 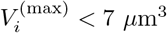 and the estimated average growth rate *G*_*i*_ *<* 10 *μ*m^3^/hour, where

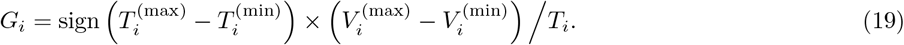

We also ignore any tracks that have no identified lineage relationships: they are neither assigned as a mother nor as a bud/daughter at any time point.

Our tracking algorithm correctly identifies many instances where a mother and bud pivot with the flow of the medium, but exceptions do arise. For a given mother, we therefore join contiguous daughter tracks – pairs of daughter tracks where one ends with the other starting on the next time point – if the extrapolated volume of the old track falls within a threshold difference of the volume of the new track. Specifically, for the pair of contiguous tracks *i* and *j*, with track *i* ending at time point *t* and track *j* beginning at time point *t* + 1, we calculate

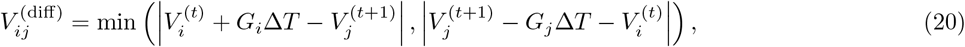

where 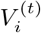 is the volume of track *i* at time point *t* and Δ*T* is the time step between time points *t* and *t* + 1. We join these tracks if 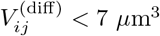.

Finally, we discard any daughter tracks with fewer than five time points.

## Appendix 5

### Estimating birth times from fluorescent markers

**Figure 1:**
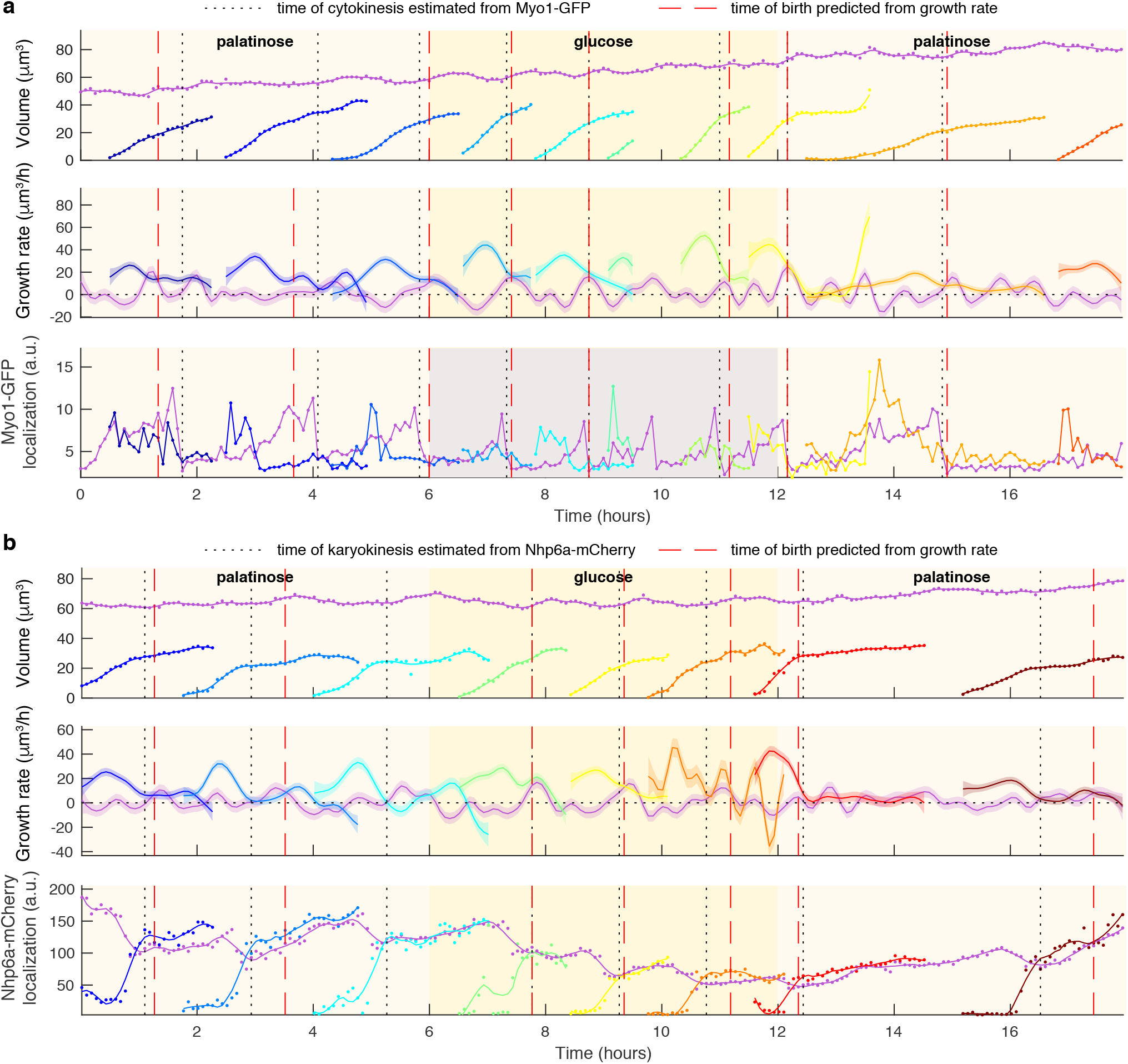
Markers for karyokinesis and cytokinesis reveal coincidence with a crossing point in mother and daughter growth rates. **a** Time series for a mother (purple) and its buds/daughters for a switch from 2% palatinose to 2% glucose and back. Volumes and growth rates are as estimated by BABY. Localization of Myo1-GFP to the bud neck is used to identify times of cytokinesis (vertical black dotted lines) as described in the Supplementary Text. For comparison, birth times predicted by our growth rate heuristic are also shown (vertical red dashed lines). **b** As for (a), but using localization of Nhp6A-mCherry to the nucleus to identify times of karyokinesis (vertical black dotted lines). Both the raw (points) and smoothed (lines; Savitzky-Golay filter with third degree polynomial and smoothing window of 15 time points) localization of Nhp6A-mCherry is shown.

We used either Myo1-GFP or Nhp6a-mCherry to estimate the time at which a bud becomes an independent daughter. Myo1 – a type II myosin – localises to the bud neck and shows a drop in intensity upon cytokinesis (Appendix 5 Fig. 1a); Nhp6a, a histone-associated protein localised to the nucleus, shows a drop in intensity upon karyokinesis (Appendix 5 Fig. 1b). Although karyokinesis and cytokinesis are distinct events, their timing is similar [24], and we assume both mark the birth of an independent daughter. For Fig. 4a & b, Appendix 5 Fig. 1a and 2, we used Myo1-GFP; for Fig. 4d & e, Fig. 5, Appendix 5 Fig. 1b, Figure 4—figure supplement 1, Figure 5—figure supplement 1, and Figure 6—figure supplement 1, we used Nhp6a-mCherry.

Cytokinesis is accompanied by a drop in Myo1-GFP intensity at the bud neck in the mother cell (Fig. 4a), and we use this phenomenon to determine when cytokinesis occurs, assuming that it is fast compared to the time interval of our imaging. We use the time series of fluorescence localisation within the mother cell over the period where the mother has a bud and estimate its time derivative using finite differences (NumPy’s `diff` function). To obtain candidate time points for cytokinesis, we find the dips in this derivative with a minimum prominence, via SciPy’s `findpeaks` function. The actual time of cytokinesis is taken to be the candidate point with the minimal derivative, corresponding to the strongest down-shift in fluorescence.

For Nhp6a-mCherry, the timing of karyokinesis is less obvious, although when the nucleus partitions between the mother and its bud there is both a fall in fluorescence localisation in the mother and a rise in the bud (Appendix 5 Fig. 1b). As karyokinesis completes, both localisation signals level to a similar, often slowly increasing value. We therefore identify the completion of karyokinesis as the time point when fluorescence localisation shifts from a fast increase in the bud and decrease in the mother to slow increase in both. We thus search for a minimum in the time derivative of the localisation signal for the mother that occurs near a maximum in the time derivative of the daughter.

Specifically, we consider a mother and bud/daughter each with a time series of fluorescence localisation. We apply a Savitzky-Golay filter with a third order polynomial and a smoothing window *W* to obtain the smoothed localisations, denoted *m*_*t*_ and *b*_*t*_, and their time derivatives. We find candidate time points of karyokinesis by identifying peaks in the series 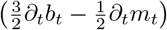 that have a minimal peak prominence *P*, using Matlab’s `findpeaks` function. For each peak, we define candidate intervals of karyokinesis as *{*|*t t*_*i*_| *?* round (*w*_*i*_*/*2)*}* where *t*_*i*_ is the time that the peak occurs and *w*_*i*_ is its width. This set of candidates is filtered by thresholding on peak height, the maximal value of the bud’s smoothed localisation, the range of the bud’s localisation, the minimal value of the mother’s localisation minus the maximal value of the bud’s, and the minimal value of the time derivative of the mother’s localisation, all calculated over the candidate time interval. The smoothing window *W*, minimal peak prominence *P*, and each threshold are manually curated for every experiment. The first occurring peak that passes all filters defines the time interval of karyokinesis, and karyokinesis’s time of completion is taken as this interval’s final time point.

### Predicting cytokinesis from growth rate

All together, we are able to determine key events of the cell cycle. First, we define a cell cycle for each mother as the duration between two birth points, obtained from the lineage assignment. These points approximately correspond to shortly after the START checkpoint [76]. Second, assuming that the births are accurately predicted, we identify a single point of cytokinesis within the corresponding cell cycle.

We observe three phases of growth during a cell cycle (Fig. 4a-b). First, the bud dominates growth during S/G2/M, with its growth rate peaking midway through that period while simultaneously the mother’s growth rate falls. Second, the bud’s growth rate decreases as cytokinesis is approached. Near cytokinesis, the mother’s and bud’s growth rate have similar magnitudes, becoming identical at multiple timepoints. Finally, the mother’s growth rate increases after cytokinesis, peaking during G1.

Using these observations, we developed an algorithm to identify the point of cytokinesis. Our growth rates are defined by both a mean and a standard deviation because they are estimated using Gaussian processes, and we use both quantities to compare growth rates of a mother and its bud. Our algorithm determines the point of cytokinesis as the first time point where the mother and bud’s growth rates are similar, within a threshold *g*, and the bud’s growth rate is sufficiently large, above a threshold *g*_*d*_. This additional constraint eliminates false negatives when the growth rate of both mother and daughter undergo a sudden drop, which may occur when the extracellular media is changed. In practice, we set *g*_*d*_ = *g*, and the threshold *g* is a parameter whose optimal value varies from cell cycle to cell cycle, especially with changing extracellular media. A too small *g* gives poor accuracy for fast-growing cells where the difference between the mother’s and bud’s growth rates is larger; a too large *g* gives poor accuracy because cytokinesis is predicted too early. We therefore choose *g* from a range *g*_0_,…, *g*_*n*_ chosen manually to increase the number of cell cycles for which cytokinesis is found, without affecting the error (Appendix 5 Algorithm 2). If the algorithm fails to find a value of *g* that gives a point of cytokinesis, the cell cycle is marked as unknown and ignored in further analysis.

We evaluated the algorithm by the correlation between the real time of cytokinesis and the predicted time (Appendix 5 Fig. 2a & c). The prediction is both accurate and robust to different media, with correlation coefficients between 0.939 and 0.998 for two different experiments and three different media. The algorithm is also usually accurate at transitions between two media, when START occurs before and ends after the transition. This result suggests that the relationship between the mother’s and bud’s growth rates is informative despite transient changes in growth rate (Fig. 4b).

**Figure 2:**
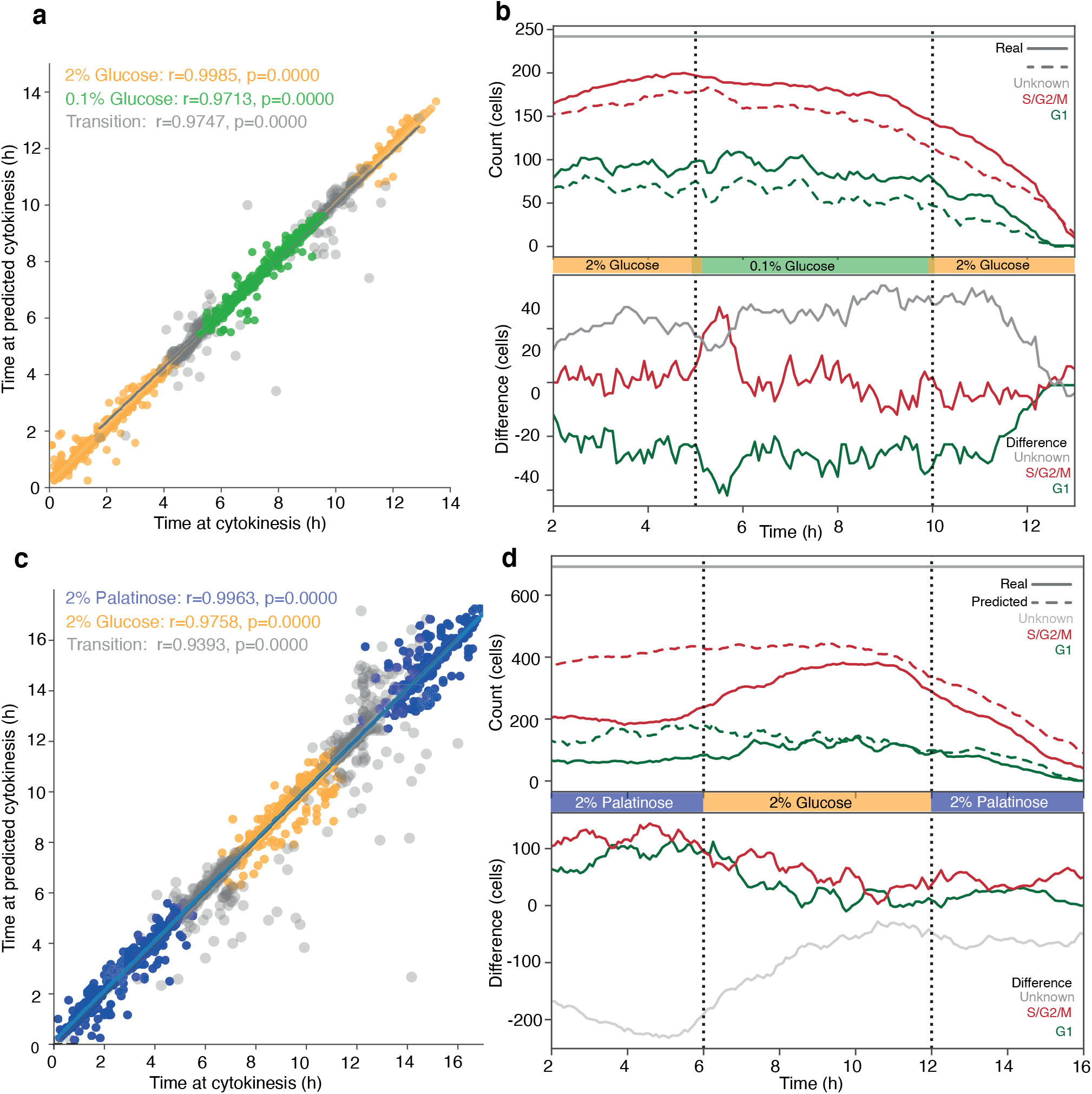
Performance of the prediction of cytokinesis from growth rate. The cells in **a** and **b** were imaged for five hours in SC media with 2% glucose, five hours in SC media with 0.1% glucose, and five hours in SC media with 2% glucose. The cells in **c** and **c** were imaged for six hours in SC media with 2% palatinose, six hours in SC media with 2% glucose, and six hours in SC media with 2% palatinose. **a, c** Comparison of the real vs predicted (absolute) time of cytokinesis for each cell cycle. The points are separated by color based on which condition the cell cycle occurs in, with cell cycles that span a switch marked as Transition (grey). **b, d** Cumulative counts of mother cells in each cell cycle stage throughout the experiment, with vertical lines representing the switch (top), with real (filled) and predicted (dashed) cytokinesis prediction used to delineate between G1 and S/G2/M. The gray line shows the total number of cells considered in the experiment, defining how many were in an unknown cell cycle state at any given time. The bottom panel shows the absolute difference between real and predicted number of cells in each cell cycle phase, as well as in unknown states, across the experiment.

Further, we verified that any prediction errors do not change the distribution of cell cycle phases over time. We used the real and predicted points of cytokinesis to split cell cycles into G1 and S/G2/M and counted how many mother cells are in each phase at any given time. We then compared these cumulative counts (Appendix 5 Fig. 2b & d). For a switch from high to low glucose, although the number of unknowns is larger for the prediction – the cumulative count of G1 and S/G2/M is smaller, the prediction still captures the dynamics of the population (Appendix 5 Fig. 2b). For a switch from palatinose into glucose, the accuracy is less (Appendix 5 Fig. 2d). The oscillations in the number of cells in G1 that occur in glucose are weaker, but we do predict cells accumulating in G1 once palatinose returns.

We note that these comparisons necessarily use only cells that are assigned at least one valid cell cycle with the Myo1 method. In the palatinose-glucose-palatinose experiment in particular, there are multiple cells that are therefore ignored despite being assigned at least one valid cell cycle with the growth-rate method. It is possible that discarding these cells affects the overall result.

#### Algorithm 2: Cytokinesis from growth rate

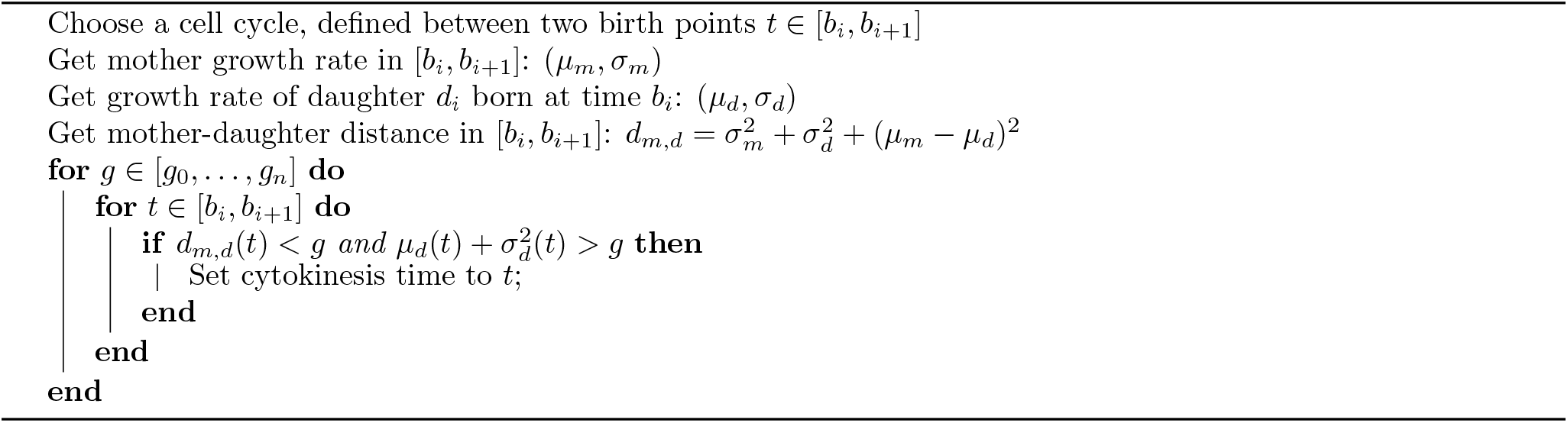

## Appendix 6

### Correlating nuclear Sfp1 with growth rate

The cross correlation of time series can reveal regulatory relationships [77]. Here we apply cross correlations to investigate if fluctuations in Sfp1’s localisation anticipate fluctuations in growth rate. Analysis by the method of Kiviet *et al*. [21] assumes steady-state cells. We nonetheless make use of data with a switch from palatinose to glucose and back (Fig. 4b), but limit to time points from either the four hours preceding the switch to glucose – approximately steady growth in palatinose – or the four hours preceding the switch back to palatinose – approximately steady growth in glucose.

Correlations may occur on scales longer than the duration of a cell cycle, so we limit our analysis to mother cells that are present over the full four hours of steady growth. We use the summed mother and bud growth rates whenever a bud is present because most of the mother’s growth is in the bud. We identify when a daughter separates from its mother using Nhp6A-mCherry (Appendix 5). Almost all daughters are washed away before they become mothers, making the lineage trees in our data unbranched and simplifying the analysis.

For each mother *i*, we have a time series 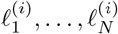 of the degree of localisation of Sfp1-GFP and a time series of instantaneous growth rates 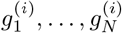. For our sampling interval of Δ*t* = 5 min, *N* is 48. We denote the total number of mother cells by *M* and calculate the deviation from the population mean for each time series:

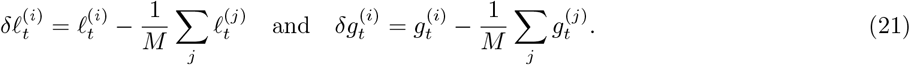

The cross-covariance of Sfp1 localisation and growth rate at a time lag of *r* Δ*t* is then [21]:

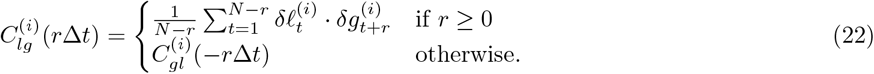

We find the cross-correlation through normalising by the standard deviations:

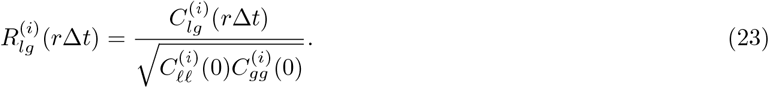

We determine the auto-correlation for Sfp1 localisation, 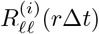, and for growth rate, 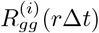, similarly. In Fig. 5c–d of the main text, we show the mean and 95% confidence interval over all mother cells (all *i*).

## Appendix 7

### Real-time feedback control

In these experiments, we wished to trigger a change in media based on the cells’ growth rate. As an example, we switched medium from a richer to a poorer carbon source and used BABY to determine how long cells should be kept in this medium if we want approximately 50% to have resumed dividing before switching back to the richer medium.

Code to implement the feedback control was run on two computers, one controlling the microscope (Appendix 7 Algorithm 3) and the other both segmenting images (via calls to Python functions) and determining growth rates (Appendix 7 Algorithm 4). The code was written in Matlab and is available on request.

We defined the fraction of escaped cells as the proportion of included mothers that have had a bud/daughter exceed a threshold in growth rate of 15*μ*l/hour at any time point after the onset of the lag in growth caused by the poorer carbon source. We define this lag period to begin at the time point when the median daughter growth rate first drops below 5*μ*l/hour. To be included a mother cell must satisfy two requirements: be present in our data for at least 95% of the time points from the 20 time points before the first switch to the current time point; and be assigned a bud/daughter for at least 10% of the time it was observed.

To increase processing speed, we use Savitzky-Golay filtering to estimate growth rates. The resulting first derivative is not well constrained at the end-points, making instantaneous growth rates vary widely at the most recently acquired time point. We therefore used growth rates up to and including the time point three steps before the most recent when determining the fraction of escaped cells.

We used the strain BY4741 Sfp1-GFP Nhp6A-mCherry in both experiments.

#### Algorithm 3: Feedback control – pseudocode for microscope acquisition software

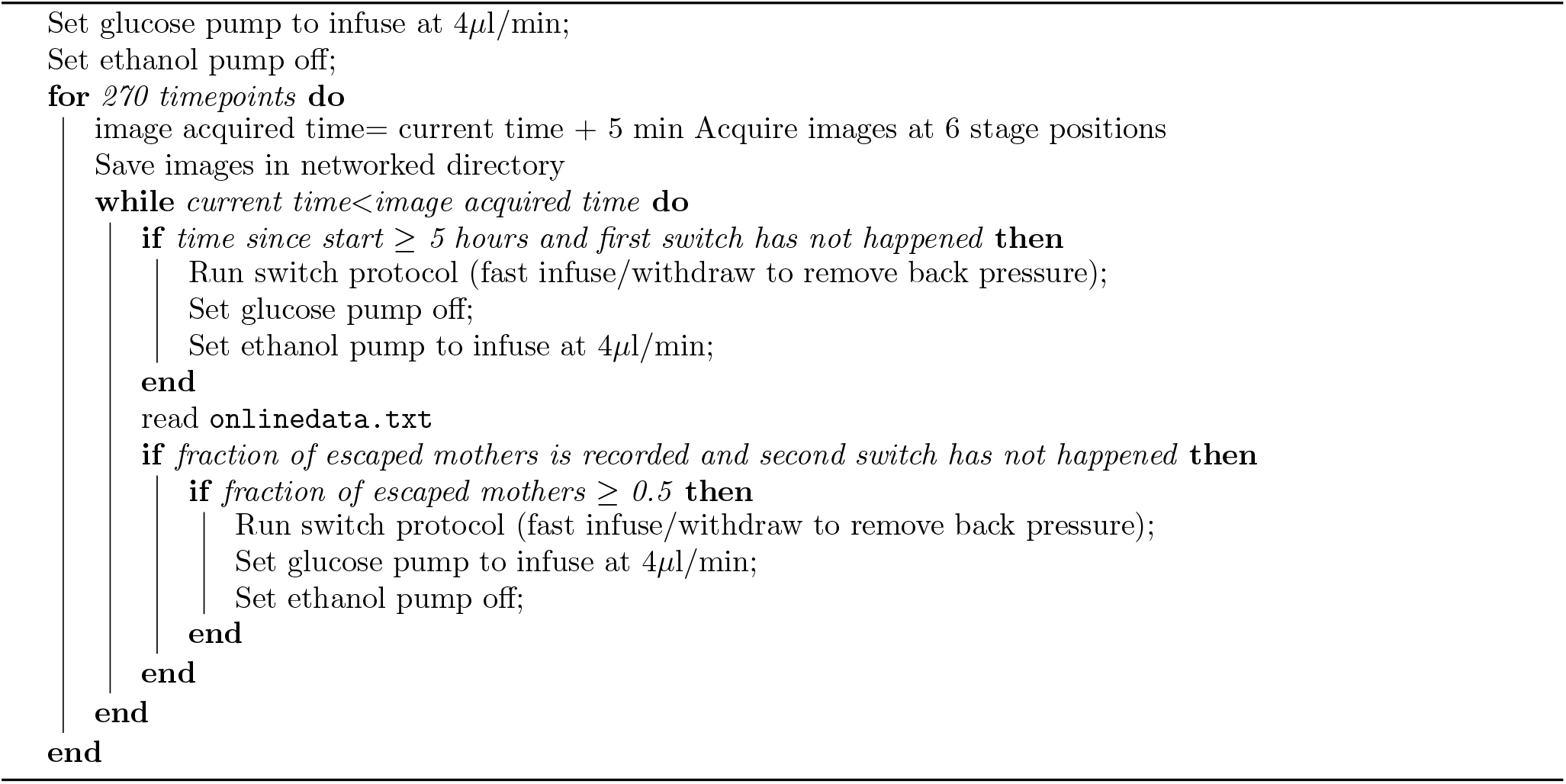

#### Algorithm 4: Feedback control – pseudocode for segmentation software

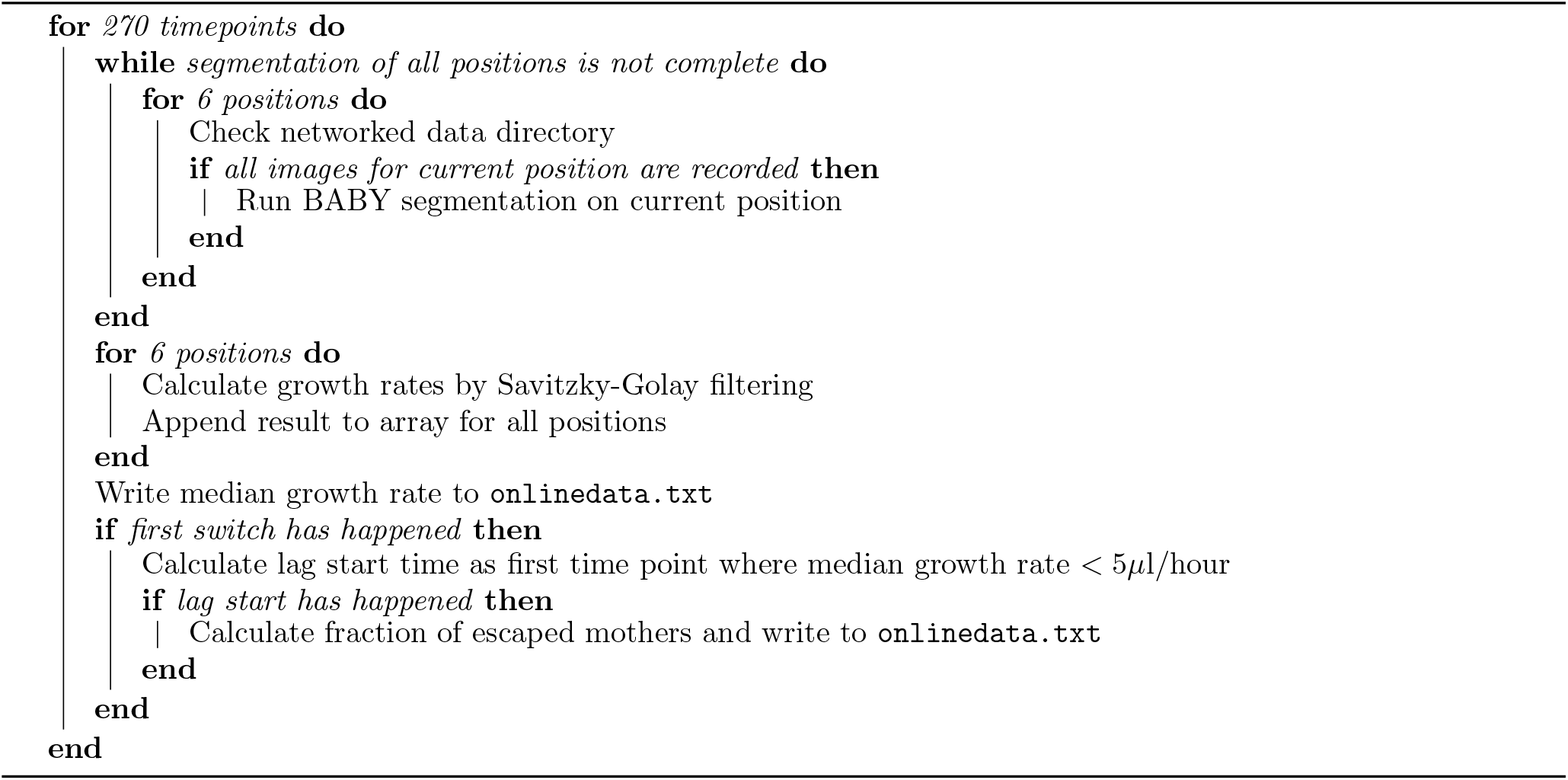

**Figure 1—figure supplement 1:**
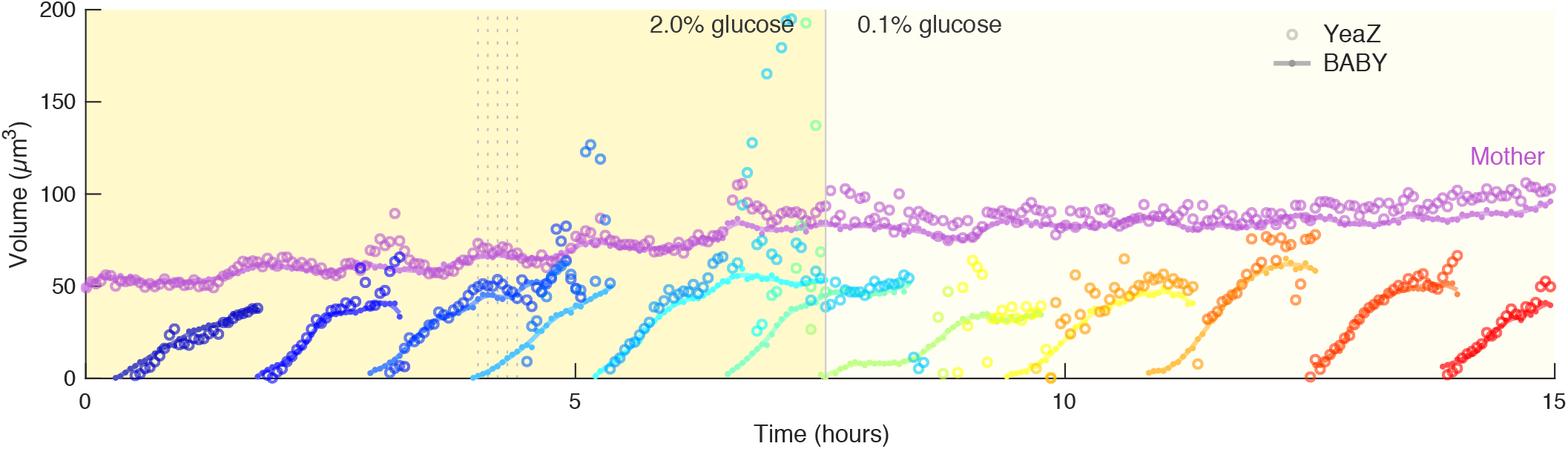
Segmentation methods without overlaps can underestimate bud area or fail to detect buds. The YeaZ segmentation algorithm [34], which segments cells from the semantic output of a CNN without allowing for overlaps, was used to segment and track the same images used to generate the time series of Fig. 1e. We show volume time series – estimated from the area of each segmented mask assuming a spherical shape – for all tracks that overlap (mask IoU *>* 0.5) with those of the segmented output of BABY (light filled circles). As YeaZ only accepts a single Z section as input, we performed a systematic search over both Z section and segmentation parameters and show the closest match to the data obtained with BABY. Vertical dashed lines correspond to the images shown in Fig. 1b.

**Figure 3—figure supplement 1:**
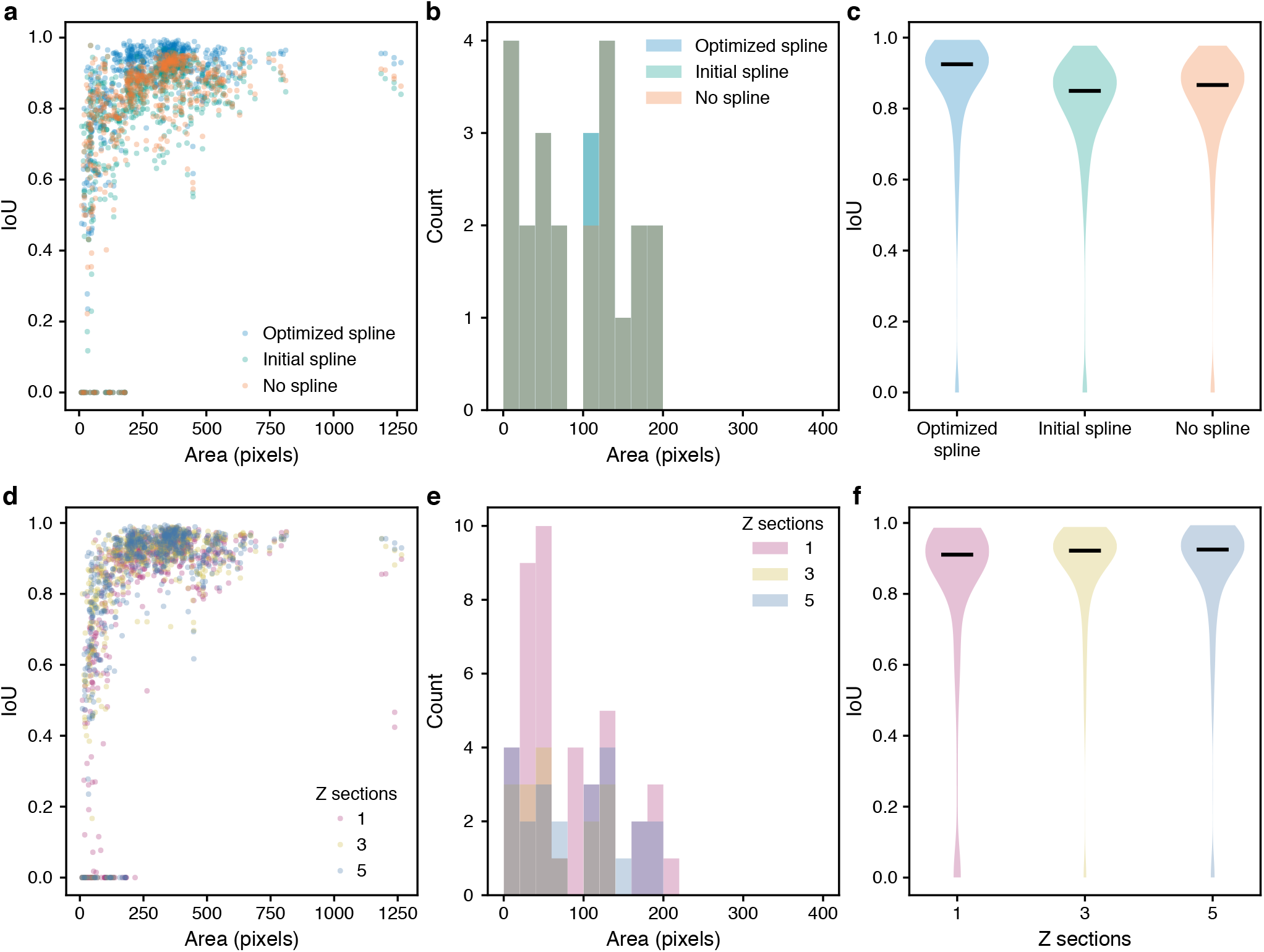
Segmentation accuracy is improved by edge optimisation and use of multiple Z sections as input. **a–c** The intersection over union (IoU) with curated masks was evaluated on test data for the predictions of selected stages of the BABY algorithm: masks predicted directly from CNN output with appropriate morphological dilation (no spline), preliminary fits of these masks with an equal-angle radial spline (initial spline), or optimised fits of the radial spline to the edge output of the CNN (optimised spline). **d–f** The IoU with curated masks was evaluated on test data for the predictions of the full BABY algorithm with CNNs trained to use either 1, 3 or 5 bright-field Z sections as input. **a, d** Distribution of IoU with curated outline area. **b, e** Distribution of curated outline area for predicted outlines with IoU of zero. **c, f** Violin plots summarising IoU distributions. Black horizontal line is the median.

**Figure 3—figure supplement 2:**
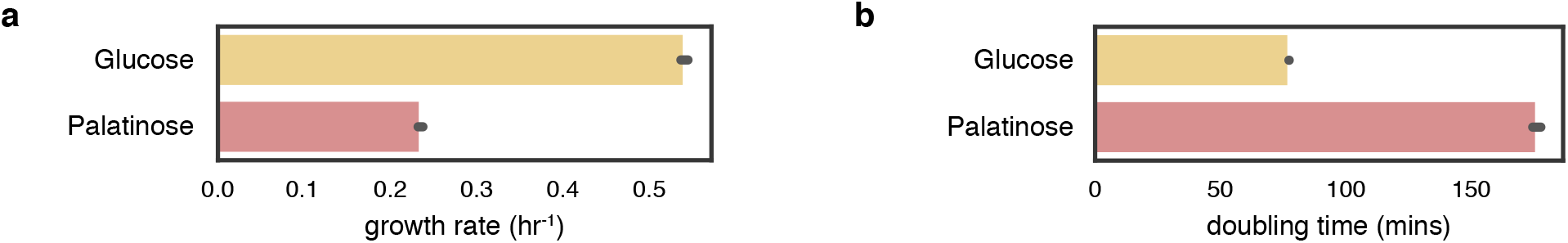
Population growth rates measured by optical density. **a** The maximal growth rate for cultures of BY4741 grown in SC with either 2% glucose or 2% palatinose. We plot the greatest local maxima in growth rate as a function of time for 47 hours of growth in a plate reader (Infinity M200, Tecan Group Ltd., Switzerland). Cells were cultured for two days in SC with 2% sodium pyruvate at 30°C, diluted 6 with fresh SC-pyruvate and grown a further 6 hours before being spun down, washed in SC with no carbon source, spun down again, resuspended in SC, added to a 96-well plate with appropriate carbon sources in each well and placed in the plate reader [78]. The time-varying growth rate was found from the optical density using a Matern Gaussian process [41]. Error bars are 95% confidence limits estimated by bootstrapping two biological replicates, each with two technical replicates. **b** Doubling times as calculated from the growth rates in **a**.

**Figure 4—figure supplement 1:**
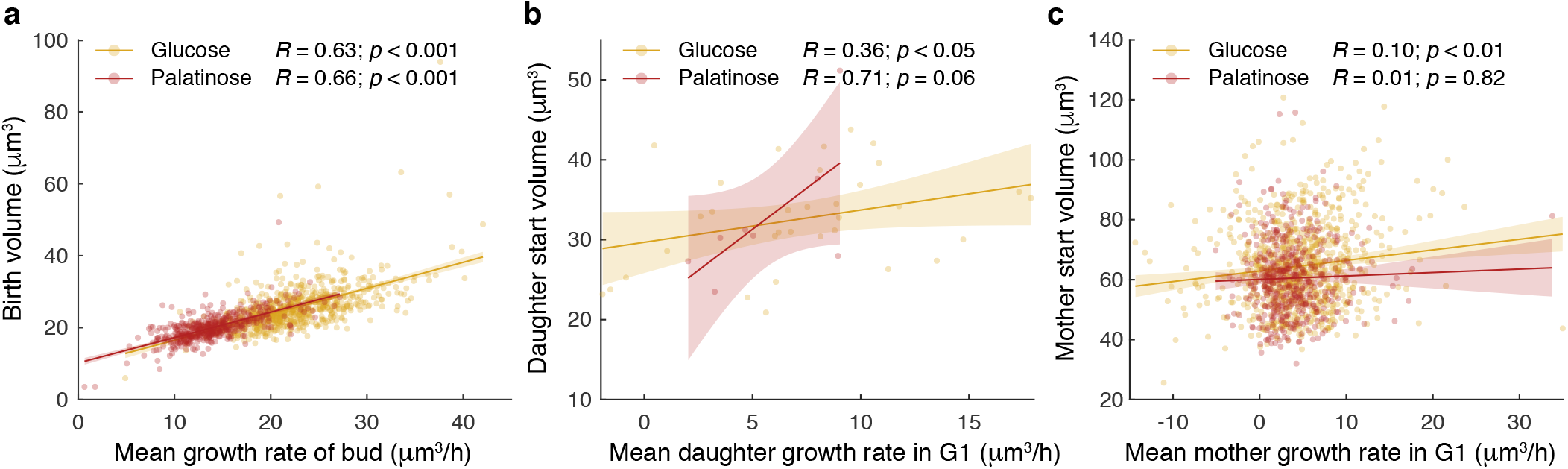
Growth rates estimated with BABY show expected correlations with volume. **a** The mean growth rate of a bud from first detection to birth (completion of karyokinesis determined from an Nhp6A-mCherry marker as described in the Supplementary Text) correlates with its volume at birth for buds growing in either 2% glucose or 2% palatinose. **b** The mean growth rate of a daughter across its first G1 phase (from birth to detection of its first bud) correlates with its volume at START (detection of its first bud). In our devices, most daughters are washed away before the appearance of their first bud. **c** In contrast, the mean growth rate of a mother during G1 (from birth of a daughter to appearance of the next bud) shows little to no correlation with its volume at START (appearance of its next bud).

**Figure 5—figure supplement 1:**
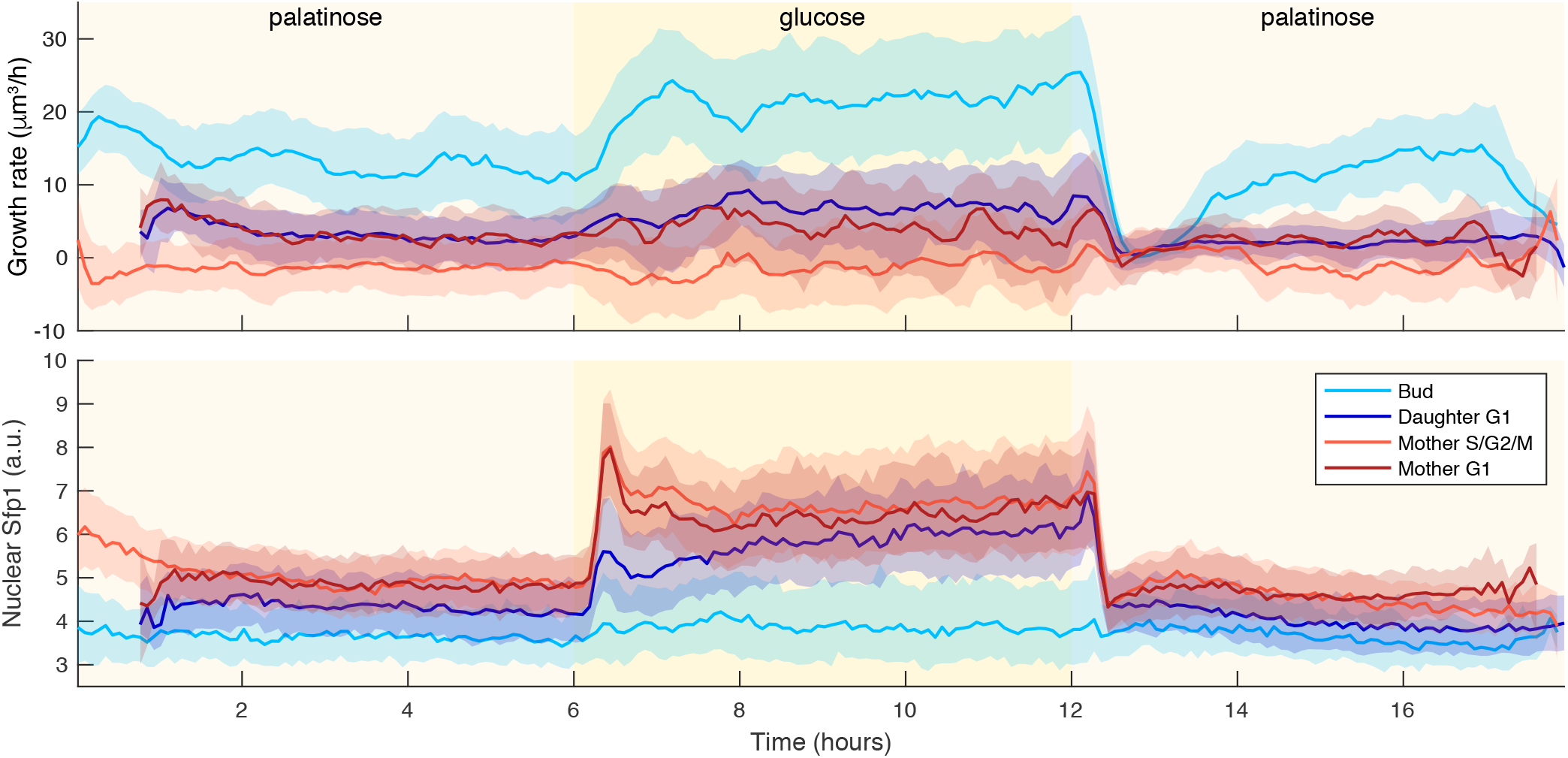
Irrespective of cell cycle phase, growth rates transiently drop for a shift to a poorer carbon source. Time series of median growth rate (upper) and median Sfp1-GFP localization (lower) across different cell cycle phases for a switch from 2% palatinose to 2% glucose and back. A Nhp6A-mCherry reporter was used to identify completion of karyokinesis (Supplementary Text) so that the time series for each cell could be partitioned into budding – detection of bud up to karyokinesis, daughter G1 – from karyokinesis up to the detection of its first bud, mother S/G2/M – from detection of a bud to karyokinesis, and mother G1 – from karyokinesis to the appearance of its next bud. The interquartile range is shaded.

**Figure 6—figure supplement 1:**
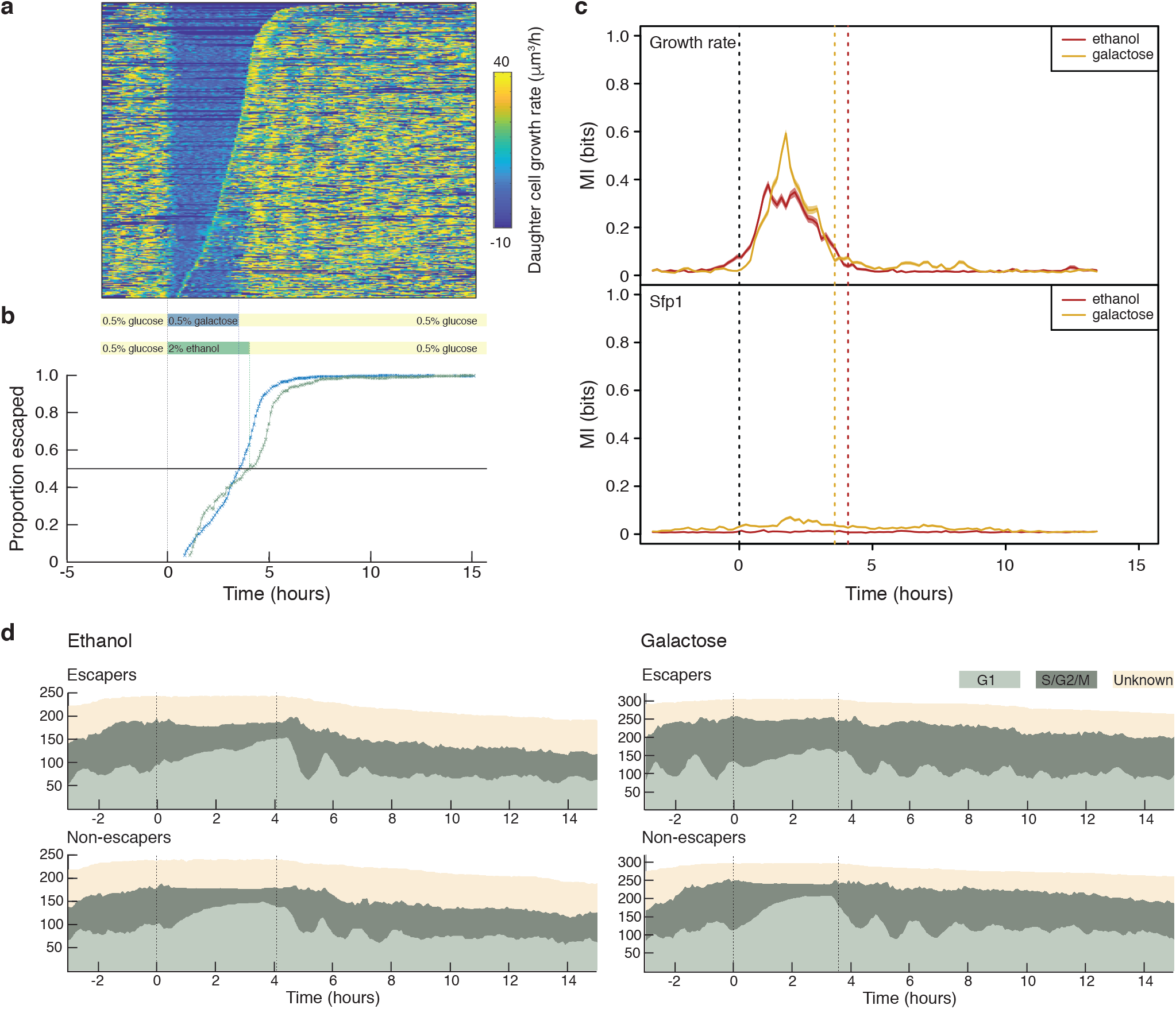
Live growth rate control of media flow. **a** Kymograph showing the daughter cell growth rates calculated during image acquisition for an experiment in which the carbon source in of the second media is 0.5% galactose. The experimental method is otherwise identical to the experiment with ethanol shown in Fig. 6 of the main text. Rows representing mother cells are sorted by the time each cell resumes division, with cells resuming later shown toward the top. Times of the media changes are indicated below. **b** Time course of the proportion of cells that have resumed division (“escaped”) following the switch to 2% ethanol (green) or 0.5% galactose. The times of the two media switches are indicated for each experiment. **c** Estimated lower bound on the mutual information between escaping/non-escaping cells and either daughter growth rate (top) or nuclear localization of Sfp1 (bottom) in experiments with ethanol or galactose as the second carbon source. Mutual information was estimated by the decoding method [20] for a sliding time-window of 100 minutes, with the mean and 95% confidence interval (shaded area) of 100 bootstraps of the estimation algorithm shown. Decoding was by a gradient boosting model trained on concatenated raw, Fourier-transformed, and rank-ordered versions of each time window. **d** Time courses of estimations of cell cycle stage for escapers and non-escapers for the experiments using ethanol (left) and galactose (right). There is an accumulation of cells in G1 during the period in the second carbon source and synchronisation in cell cycles is evident from the periodic increases and decreases of the proportions of cells in G1 following the reintroduction of glucose at the second media switch.

## Notes

### Competing Interest Statement

The authors have declared no competing interest.

